# Evidence for the pairing of mRNA exiting ribosomes acting as a Driver of unidirectional forward bypassing: Its cessation permits Backwards scanning

**DOI:** 10.1101/2024.12.09.627546

**Authors:** Norma M. Wills, Krishna Parsawar, John F. Atkins

**Affiliations:** Department of Human Genetics, University of Utah, Salt Lake City, UT 84112, USA; Mass Spectrometry and Proteomics Core Facility, University of Utah, Salt Lake City, UT, USA; Schools of Biochemistry and Microbiology, University College Cork, Cork T12XF62, Ireland

**Keywords:** Translational bypassing, T4 gene *60*, frameshifting, decoding, ribosome versatility

## Abstract

Efficient translational avoidance of a 50 nucleotide non-coding insert in phage T4 gene *60* sequence involves dissociation of peptidyl-tRNA pairing from the mRNA ‘take-off’ codon, GGA. This tRNA is retained in the ribosomal P-site during bypassing. Its anticodon does not scan the 5’ part of the non-coding sequence, which contains a cognate GGG codon, but it does scan the main and 3’ part of the coding gap for potential complementarity. Re-pairing to a matched GGA ‘landing site’ codon occurs exclusively 48-50 nt after take-off. Coding resumption ensues. Here we analyze proteins synthesized from mutants that abrogate the potential for re-pairing at the canonical landing site. The results show that some ribosomes move backwards 39 nt allowing re-pairing to the initially hidden GGG or forwards at least 56 additional nt before resumed anticodon: codon pairing at a downstream matched codon. We discuss the nature of the driving force for the initial unidirectional movement and why it becomes dissipated when the site of landing in WT is reached. Also presented is an analysis of the relative importance of recoding signals located 6 nucleotides upstream (5’) and close downstream (3’) of the WT landing site, for re-pairing to mRNA and coding resumption.

## INTRODUCTION

Chromosomal integration of mobile elements can have a variety of consequences. Among these are Alzheimer’s relevant-retrotransposon-derived Arc that mediates tau transmission via extracellular vesicles (1), or lethal consequences due to inactivation of an essential gene. However, in decoding phage T4 gene *60*, the lethal consequences of a stop codon-containing insert associated with a homing endonuclease as part of a mobile DNA cassette (2), are avoided by efficient translational bypassing (3,4). The insert creates a 50nt coding gap that separates an ORF1 from ORF2 of a topoisomerase subunit encoding gene (3). Discontiguous translation of the two separate ORFs to synthesize a single protein involves a productive exception to a central tenant of decoding: co-linearity of mRNA and protein sequence. An exploration of counterparts in other phages has raised the possibility of a regulatory function (5). [As some topoisomerases act on RNA as well as DNA, were T4 topoisomerase to inhibit a feature associated with the distinctive ribosome conformation/mRNA structures, it would be quite extraordinary. T4 gene *60* bypassing has a strong temperature dependence because of the RNA structures involved (6–8). A study of a different case of anticodon dissociation and re-pairing to mRNA (9) is also relevant to this point.] A single bypassing event is involved in gene *60* decoding, unlike strong candidates at two positions in decoding a phage terminase gene (10,11) and the multiple bypassing events in several different mitochondrial mRNAs of *Magnusiomyces* yeasts (12–14). However, studies of gene *60* have been foundational for in-depth understanding of one type of bypassing and an indicator of how bypassing can reveal unappreciated aspects of decoding versatility.

The 50 nt coding gap in T4 gene *60* is between codons 46 and 47. Codon 46, GGA, is the bypassing ‘take-off’codon, from which the anticodon of peptidyl-tRNA dissociates at the initiation of bypassing. The peptidyl-tRNA is retained within the ribosome as it traverses along the mRNA to the matched GGA, the “landing site” codon, to which the peptidyl-tRNA re-pairs to mRNA. As this pairing occurs in the ribosomal P-site, the adjacent A-site is the resume site to which entering aminoacyl-tRNA pairs leading to a resumption of protein synthesis (4) (Fig. 1).

**Fig. 1.**
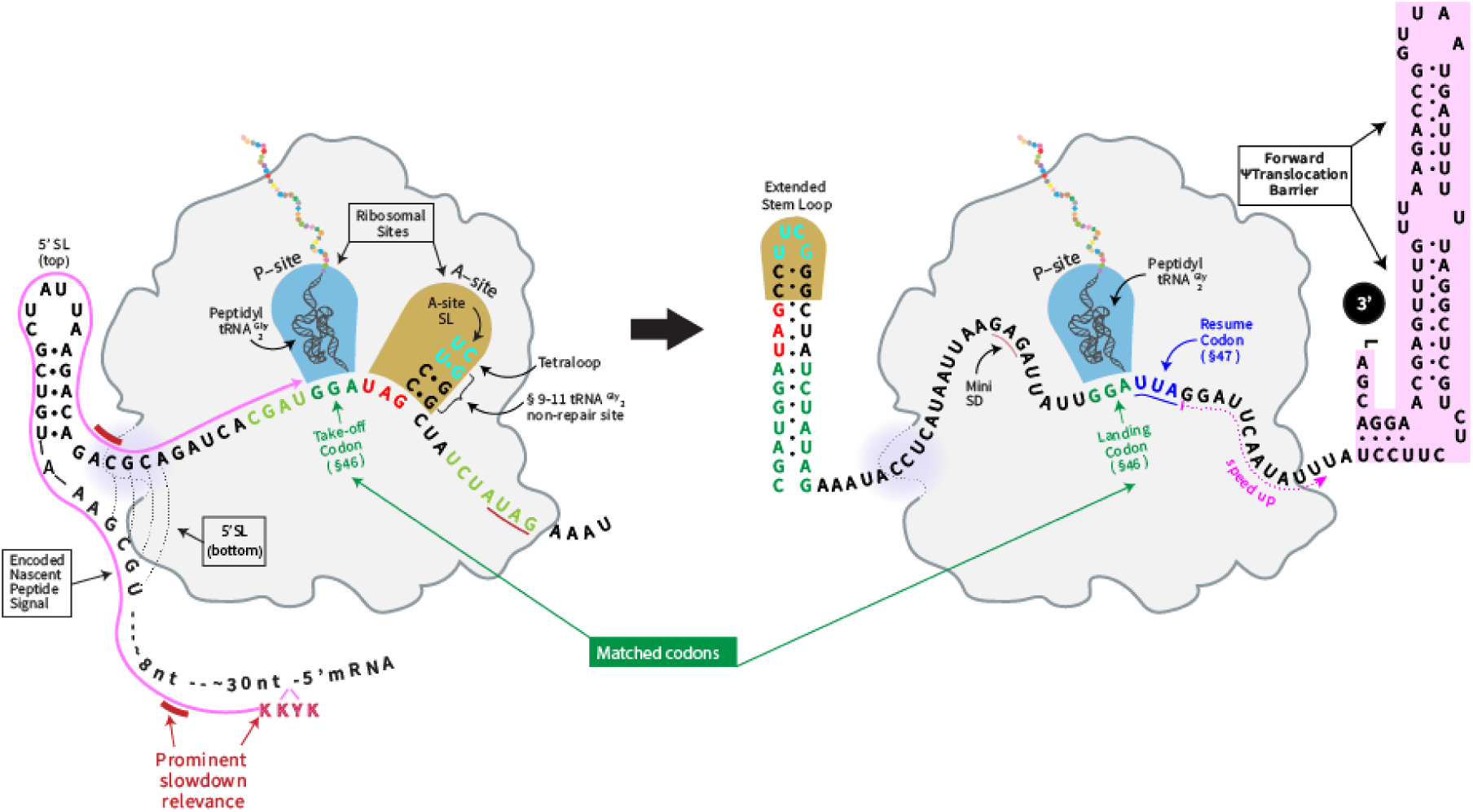
Bacteriophage T4 gene *60* translational bypassing. A compilation of the key regulatory features involved in gene *60* bypassing including: [1] the peptidyl-tRNA^Gly^ and matched take-off and landing GGA codons (dark green); a GGG at which landing does not occur in WT is right-bracketed, [2] the UAG stop codon immediately 3’ to the take-off codon (bright red), another UAG is underlined in dark red and two UAAs further 3’ in the gap, UAAuUAA, are not highlighted, [3] an upstream nascent peptide signal (KKYK-LQNNVRRSIKSSS^13-29^) [4] the 5’ stem loop, 9 nts 5’ of the take-off codon, [5] Left panel: the small mRNA ‘A-site’ stem loop that forms within ribosome, [6] Right panel: the extra nucleotides (light green) in the ‘Extended stem loop’ proposed to form on mRNA exit from the ribosome, [7] a mini SD (Shine–Dalgarno-like) sequence GAG 6 nts 5’ of the landing codon (underlined), [8] the translational resume codon UUA (blue), [9] The prominent positions of ribosome speed reduction in a progressive slow-down on approach to the take-off codon are illustrated in 3 red blocks. The nascent peptide is in pink with its KKYK sequence that interacts with the ribosome peptide exit tunnel being highlighted. Speed-up is also indicated (pink underline), [10] the Forward ψTranslocation Barrier (3’SL), 14 nts 3’ of the landing site where the anticodon re-pairs to mRNA, is also illustrated.

As shown *in vivo* (15,16) and in one *in vitro* study (8), as the anticodon of the peptidyl-tRNA^Gly^ involved in the bypassing (17) dissociates from the take-off codon GGA with essentially 100% efficiency (as elaborated on below). Ribosomes traverse the 50nt coding gap by what has been termed Pseudotranslocation (here written ψTranslocation) with a proportion dropping off before the landing site (18–20). This rate is ~1% per nt (8), consistent with the proportion of ribosomes that successfully bypass the coding gap being as high as 50% at 37° (8,15,21,22). The anticodon of the peptidyl-tRNA retained in the ribosomal P-site pairs with the GGA 48-50 nt 3’ of the take-off codon (4). The ribosome is a rotated state unusual for entry and pairing of cognate tRNA to the adjacent A-site (‘resume codon’) and associated with a substantial delay before coding resumes (8,21).

The sequence of the nascent peptide encoded 5’ of the take-off site is very important for bypassing (4). Its 9-fold effect on efficiency (18) is due to its several roles, including importantly that of peptidyl-tRNA anticodon dissociation from the take-off codon at the initiation of bypassing (6–8,15, 21, 23). Ribosomes initiating bypassing have a non-canonical hyper-rotated state (6–8,21). [Though the nascent peptide and the A-stem loop can have separate effects on bypassing, their joint effects are relevant to formation of the hyper-rotated state (8,21).] In addition to the major role of the nascent peptide: exit tunnel interaction in anticodon: codon dissociation at take-off, it has an effect on landing fidelity (18). Genetic studies have provided evidence for nascent peptide mediated effects continuing during bypassing to have some destabilizing effect on anticodon: codon interaction during the sampling of multiple successive potential codons by the single tRNA retained in the ribosomal P-site. Another role for the nascent peptide: exit tunnel interaction has emerged from recent insights into prominent slow-down positions in the progressive reduction in translation speed before the mRNA take-off site reaches the ribosomal P-site. An initial smFRET study revealed the slow-down (21) as well as the importance of a KKYK motif that was also identified in separate mutagenesis studies that showed the tyrosine was its most important component (19,24). Fig. 1. However, it took a more recent kinetics study, performed at close to the physiological speed involving take-off, bypassing and landing taking 30-40 seconds (8), to reveal the location in the exit tunnel at which the KKYK nascent peptide exerts its (at least main) effect. This study showed that translation slow-down commences ~21 codons after translation initiation, i.e ~26 codons 5’ of the take-off site. This is independent of decoding and translocation and occurs when the KKYK motif reaches the ribosome’s peptide exit tunnel’s narrowest point, the constriction site which is lined with positively charged arginine residues. [Deletion of the signaling loop of one of the two proteins, RpsL23, at the constriction site in the largely rRNA interior surface (‘wall’) of the exit tunnel reduced bypassing by 50% (19).] Electrostatic repulsion induced by positive charges in the vicinity of the constriction may contribute to the slowdown via a long-distance effect on the peptidyl transferase center of the ribosome (25). This slow-down is important for ribosome spacing reasons between successive ribosomes under polysome conditions (8). Reduced speed at this position by the second or following, ribosome in a pair of translating ribosomes would allow mRNA exiting the lead ribosome to form an intra-mRNA stem loop known as the 5’ stem loop that influences bypassing by the lead ribosome.

There are two more prominent slow-down positions though it is doubtful if the slow down at these positions have relevance for bypassing rather than being incidental consequences of features that are otherwise significant for bypassing (8). Regardless of that, at least the second of the two has significance for a possible extension of the results here involving a further enlargement of the Extended Stem Loop. The first of these occurs after translation has continued for approximately 10 more codons (Fig. 1 labeled red bar). The ribosome encounters the 5’ stem loop at its leading edge or rather just inside its mRNA entrance channel where its helicase unwinds entering mRNA structure. Then after the ribosome translates a further ~10 codons its leading edge/helicase center encounters another mRNA structure the Extended Stem Loop (Fig.1 unlabeled red bar). This is associated with a further prominent slow-down of translation rate (8).

When the take-off site reaches the ribosomal P-site, interaction of poorly understood features of the nascent chain with the interior of the exit tunnel (which is mostly rRNA), possibly even augmented by further N-terminal sequence at or after, ribosome exit, are highly likely significant for retention of the peptidyl-tRNA within the ribosome during bypassing (8,19). However, *in vivo* experiments suggest that a consequence of the nascent peptide interactions may also have the opposite effect (18). Whether, with hindsight, this is due the contributory role of the nascent peptide interactions on formation of the unusual conformation of bypassing ribosomes is unknown. However, such a trade-off could be relevant to the upper level of bypassing efficiency. Because of the likely ribosome conformation during their movement on the coding gap, their associated EF-G might be expected to be linked with recruitment of the GTP-utilizing ribosome (splitting) recycling factor, Ribosome Release Factor (RRF). If so, it could contribute to drop off. Though *in vivo* work showed that RRF does contribute to drop-off, its effect is very modest (18). That work permitted the strong deduction that the major reason for bypassing only being 50% efficient despite the ~100% take-off efficiency was due to spontaneous peptidyl-tRNA drop-off rather than other causes such as aberrant landing (18). *in vitro* studies yielded a more refined and direct conclusion about the major importance of spontaneous drop-off for bypassing efficiency (8,21). [Drop-off of peptidyl-tRNA from bypassing ribosomes, followed by Peptidyl-tRNA hydrolase (Pth) activity, liberates products identical to what, if it were to occur, would be the product derived from termination at the UAG immediately 3’ of the take-off site.]

When the take-off codon is in the P-site, there is a UAG(A) stop in the A-site where, as shown by a variety of experiments, its slow-to-decode property is important for initiation of bypassing (15). At the initiation of bypassing a small tetraloop ‘capped’ mRNA structure forms in the ribosomal A-site (6, 21). [mRNA structures in the ribosomal A-site are also known from work in a model system (26) and elsewhere (27 and refs therein).] Formation of the mRNA A-site SL contributes, at least via its role with the nascent peptide signal, to the ribosome conformation that inhibits access of release factor 1 to the UAG and so avoidance of termination (6,8,15,21). On its own the flanking sequence context of the UAG would lessen the efficiency of termination making the UAG prone to readthrough. However, in gene *60* readthrough does not occur (21,23). Access of read-through causing near cognate aminoacyl-tRNAs is also blocked by the ribosome conformation at take-off (6,8). Optimization of A-site stem loop formation contributed to irrelevance of the UAG’s poor termination context.

The A-site stem loop is also important for inactivating the effect of ribosomal protein L9 in restraining forward mRNA slippage (23) and for its tRNA-mimic role in facilitating EF-G mediated GTP hydrolysis at the initiation of bypassing (7). How nucleotides that formed the A-site stem loop pass through the ribosomal P-site is unknown, but if unfolded it is likely that part or all of them are not in the normal mRNA ‘groove’. The anticodon of the retained peptidyl-tRNA^Gly^ does not scan this 5’ region of the coding gap mRNA and so does not pair with the complementary GGG located 9-11 nts 3’ of the take-off site (21,28). However, the peptidyl-tRNA anticodon does scan the main and 3’ part of the coding gap (21). In bypassing the whole 50nt coding gap, ribosomes utilize ~90 GTPs (7). Lack of pairing of peptidyl-tRNA anticodon with this GGG during WT bypassing is pertinent to the experiments reported here. [From the 3’ end of the A-site stem loop to the landing site there are 37 overlapping triplets. Where scanning of the overlapping mRNA triplets by the peptidyl-tRNA anticodon starts is unknown, but speculatively if the scanning of each of the 37 triplets occurs and utilizes 2 GTPs as in standard translation where a single codon is sampled by multiple tRNAs, that would leave 16 GTPs for dealing with A-site stem loop nucleotides.] After mRNA sequence that formed the A-site stem exits the ribosome, it becomes part of the larger ‘Extended Stem Loop’ that is capped by the same tetraloop that capped the A-site stem loop.

In addition to the consequences of the nascent peptide just considered contributing to the fidelity of landing at the canonical site, other features enhance peptidyl-tRNA re-pairing to mRNA at the ‘correct’ landing site. One is a mini-Shine Dalgarno (SD) sequence, GAG, 6 nt 5’ of the WT landing site. Pairing of the anti-SD sequence of bypassing ribosomes with it has a restraining influence on ribosome movement past the landing site (28). This feature is conserved in a variety of newly discovered, distinct, occurrences of gene *60* bypassing where it is also embedded/ followed by an AU-rich sequence (5).

Here we analyze *in vivo*, the relative importance of these auxiliary features that enhance landing at the designated landing site in WT sequence. We also study the proteins synthesized from mutant sequence that permit deductions of ribosome movement when the peptidyl-tRNA in bypassing ribosomes cannot re-pair to mRNA at the canonical landing site. This is relevant to the function of intra-mRNA stem loops, the 5’ stem loop, and the Extended Stem Loop, both of which form as their mRNA exits the ribosome. The latter has more functionally significant base pairs (28) than what was termed the ‘take-off’ stem loop structure in the original mechanistic analysis of programmed bypassing (4). That original analysis followed studies that revealed low-level short distance bypassing/ stop hopping in the absence of stimulatory signals, i.e. where 9 nucleotides specified one amino acid; enhanced efficiency by a mutation of the relevant tRNA confirming that the phenomenon was at the translation level (29,30).

The present work involves analysis of samples of multiple protein products differing by a very small number of internal amino acids. For a previous similar study, we created a mixture of slightly distinguishable synthetic proteins in different known relative amounts prior to purification and ESI mass spectrometry. The ratio of peak heights in the mass spectra reflected the ratio of products in the mixed samples (31). This provides considerable confidence in assessing significance of relative peak heights of closely related products in the present work.

## Materials and Methods

### Construction of plasmids containing mutant gene *60* sequences

The vector used for the construction of the Nus A–gene *60* fusions was a derivative pNTH-2 (28) containing sequences encoding 6X histidine tags in all three reading frames downstream of gene *60* sequences. For the modified WT (mWT) construct, Tm1, the −1 frame TAA stop codon §36-38 in WT was eliminated by mutation of A §38 to T and T §43 to A. (This was done to retain base pairing potential between positions §38 and §43 as occurs in WT and whose importance was unknown.) Because of the presence of stop codons in the 0 and +1 frames within the gene *60* sequence, protein products could be monitored only for −1 frame events (the same frame utilized in productive WT bypassing.) In addition, sequences encoding a Strep tag were introduced between NusA and gene *60* sequences in order to potentially utilize another affinity tag for purification of products. The Strep tag was present in constructs Tm1, Tm2 and Tm4 accounting for the higher overall masses of their products in Table S1 and ESI chromatograms. The Strep tag was not included in the remaining constructs. Construct Tm10 containing CCA §48-50 at the position of the WT landing site, also lacked 9 nts encoding three amino acids, LAV, near the 5’ end of the gene *60* sequence to allow direct comparison of bypassing products with other constructs. (These comparisons were not undertaken in this study.) Sequence X was substituted for part of the Forward ψTranslocation Barrier sequence as follows: 5’…GTAATTG**CTGCTGCGAT**TTT**TTAGTTCCT**CGAAT..3’ to disrupt potential base pairing in two of the stems (substitutions in blue bold letters; see also Fig. 1). The mutant gene *60* sequences were synthesized as gBlocks (IDT) or cloned into pUC57 by GenScript flanked by SpeI and KpnI restriction sites. The SpeI-KpnI DNA fragments were ligated with SpeI-KpnI-digested pNTH-2 and transformed into BL21(DE3) STAR competent cells (Invitrogen). Individual clones were isolated and confirmed by ABI sequencing.

### Selection of mutants of the 3’ coding gap

A set of oligonucleotides was synthesized, with mixtures of bases for positions §33-47 within the coding gap. To avoid generating stop codons, base T was not provided at positions corresponding to the first nucleotide of −1 frame codons. These were used for PCR to generate gene 6*0* fragments containing CCA §48-50 in place of GGA §48-50 though all constructs had GGA at the take-off site. These were fused to *lacZ* sequence such that bypassing to the −1 frame would lead to expression of *lacZ*. As reported elsewhere (32), colonies were screened on indicator plates for enhanced β-gal activity. One mutant sequence, Tm10, was cloned into the pNTH-2 derivative and analyzed in the present study.

### Expression of NusA-gene *60* fusion protein(s)

Cultures were grown as previously described (31). Harvested cells were lysed using BugBuster (Novagen). NusA–Strep tag–gene *60*–6 × HIS proteins were purified by sequential chromatography on Ni-NTA agarose (Qiagen) and S-protein agarose (Novagen). Proteins lacking the Strep tag were purified on Ni-NTA agarose only. Eluted proteins were concentrated and washed twice with dH_2_O using Centricon 50 (Millipore) filtration devices.

### Intact Protein Analysis

Mass spectrometry analyses of intact proteins were performed on a maXis II ETD mass spectrometer equipped with an Apollo II ion funnel ESI source (Bruker Daltonics, Bremen, Germany). The source was operated in the positive (ESI+) mode at 200°C with N2 as the nebulizer and the source temperature was kept at 150°C. The capillary voltage was maintained at 4.5 kV. Ultrapure nitrogen was used as desolvation gas with a flow of 5 L/min. Ultra-high pure nitrogen was used as the collision gas. Intact protein separation and desalting was achieved following gradient elution of the intact protein from a C4 RP column (ACQUITY UPLC M-Class Protein BEH C4 Column, 300Å, 1.7 µm, 300 µm X 50 mm). The gradient consisted of 5 to 80% solvent B in 78 mins (solvent B: 100% acetonitrile with 0.2% formic acid; solvent A: 100% water with 0.2% formic acid) was used for the analyses. Spectra were processed and then deconvoluted into molecular-mass spectra (i.e. processed into neutral molecular weight) using Data Analysis software with MaxEnt (Bruker Daltonics, Bremen, Germany).

### MS/MS of proteolytic products

In-gel digestion was performed on the Bypass products separated by SDS-PAGE. Samples were run on 15% acrylamide gels and stained with Coomassie Brilliant Blue. The proteins in the expected size range were excised and then de-stained twice in 1 ml volumes of 50% methanol with 50 mM ammonium bicarbonate (AMBIC) at room temperature while being gently vortexed for about 1 hr. The gel slices/spots were re-hydrated in 1 ml of 50 mM NH_4_HCO_3_ for 30 mins at room temperature then the gel bands/spots were cut into several pieces. Proteins were reduced by 10 mM DTT at 60°C for 30 mins and alkylated by 55 mM iodoacetamide (IAA) at room temperature in the dark for 30 mins. Gel pieces were washed with 50 mM AMBIC, dehydrated with ACN, and dried using a SpeedVac. Proteolytic digestion was performed with 20 ng/μl trypsin and Lys-C mix dissolved in 50 mM AMBIC and was incubated at 37°C overnight. Alternatively, chymotrypsin digestion was incubated at 37°C for 3-4 hrs. The digestion was quenched by the addition of 20 μl of 1% formic acid. This solution was allowed to stand and peptides that dissolved in the 1% formic solution were extracted and collected. Further extraction of peptides from the gel material was performed twice by the addition of 50% acetonitrile with 0.1% formic acid and sonicated at 37°C for 20 mins, and these solutions were also collected and combined. A final complete dehydration of the gel pieces was accomplished by addition of 20 μl of 100% acetonitrile and incubation at 37°C for 20 mins. The combined supernatant solutions of extracted peptides were combined and dried in a vacuum centrifuge (Speed-Vac). The peptides were reconstituted in 100 μL of 5% acetonitrile with 0.1% formic acid for LC/MS/MS analysis.

### LC/MS/MS Analysis of Peptides

Digested peptides were analyzed using a nano-LC/MS/MS system equipped with a nano-HPLC pump (2D-ultra, Eksigent) and a maXis II ETD mass spectrometer (Bruker Daltonics, Bremen, Germany). The maXis II ETD mass spectrometer was equipped with a captive spray ion source. Approximately 5 μl of peptide samples were injected onto a dC18 nanobore LC column. The nanobore column was made in house using dC18 (Atlantis, Waters Corp); 3 μm particle; column: 100 μm ID x 100 mm length. A linear gradient LC profile was used to separate and eluted peptides with a constant total flow rate of 400 nL/min. The gradient consisted of 5 to 96% solvent B in 78 mins (solvent B: 100% acetonitrile with 0.1% formic acid; solvent A: 100% water with 0.1% formic acid) was used for the analyses. Primary mass spectra are acquired in the time of flight (ToF) part of the instrument. Peptide molecular masses were measured by ToF yielding primary mass spectra of peptides with mass errors typically less than 3 ppm. Peptide sequencing was performed by collision-induced dissociation (CID) in the quadrupole part and measure in the ToF part of this hybrid instrument yielding fragment ions with mass errors typically less than 3 ppm as well.

### Protein ID and Database Searches

All identified peptides from protein digests were assigned from protein database searches, using in-house processing with MASCOT search engine (in-house licensed, ver. 2.5, Matrix Science, Inc.). Mascot searching was performed from an in-house computer, in which NCBI or “custom” protein databases are searched. Peptides from enzymatic digests (e.g. trypsin or chymotrypsin) were searched with enzyme-specific cut sites and identified based on the “MS/MS” Mascot search option, with the following criteria: [1] Accurate mass measurement of peptide molecular ions by ToF were performed with an 11 ppm search window (though peptides typically have less than 3 ppm mass error). Molecular ions with +2, +3, or +4 charge states determined from a ToF primary mass spectra were usually considered. [2] Peptide sequence information from MS/MS; CID fragmentation of the parent ion of each peptide was obtained in the quadrupole region of the maXis II ETD mass spectrometer. 11 ppm mass error tolerance was allowed for peptide fragment ion masses in the search (though MS/MS fragment ions typically have errors less than 3 ppm). [3] Mass data peak lists for the Mascot searches were generated using Data Analysis software (Bruker Daltonics, Bremen, Germany). [4] Peptide modifications were included in the search (e.g. oxidation on methionine; carbamidomethylation on cysteine amino acid). [5] Searches were typically performed for trypsin-specific peptide cleavages, but other cut site specifications were performed depending on the protease used. “Non-specific” cleavages were also searched. Two missed cleavages were allowed. [6] All identified peptides showed MASCOT scores greater than 20. Mascot threshold cutoffs for acceptable identified peptides typically have MASCOT scores > 20, mass errors < 3 ppm, and values less than 1ppm was expected. These cut-offs were used, with other criteria being available for checking peptide assignment.

## RESULTS

### Mass spec analysis of the effects of predetermined changes 3’ of the 5’ stem loop on landing site selection

Gene *60* sequences were cloned into vector pNTH-2 (28). The derivative proteins were purified based on C-terminal His-tags encoded in all 3 reading frames (Fig.2 panel A). Consequently, products derived from termination at internal positions were not included in the protein preps analyzed by mass spectrometry to determine landing site(s) utilized during synthesis.

**Fig. 2.**
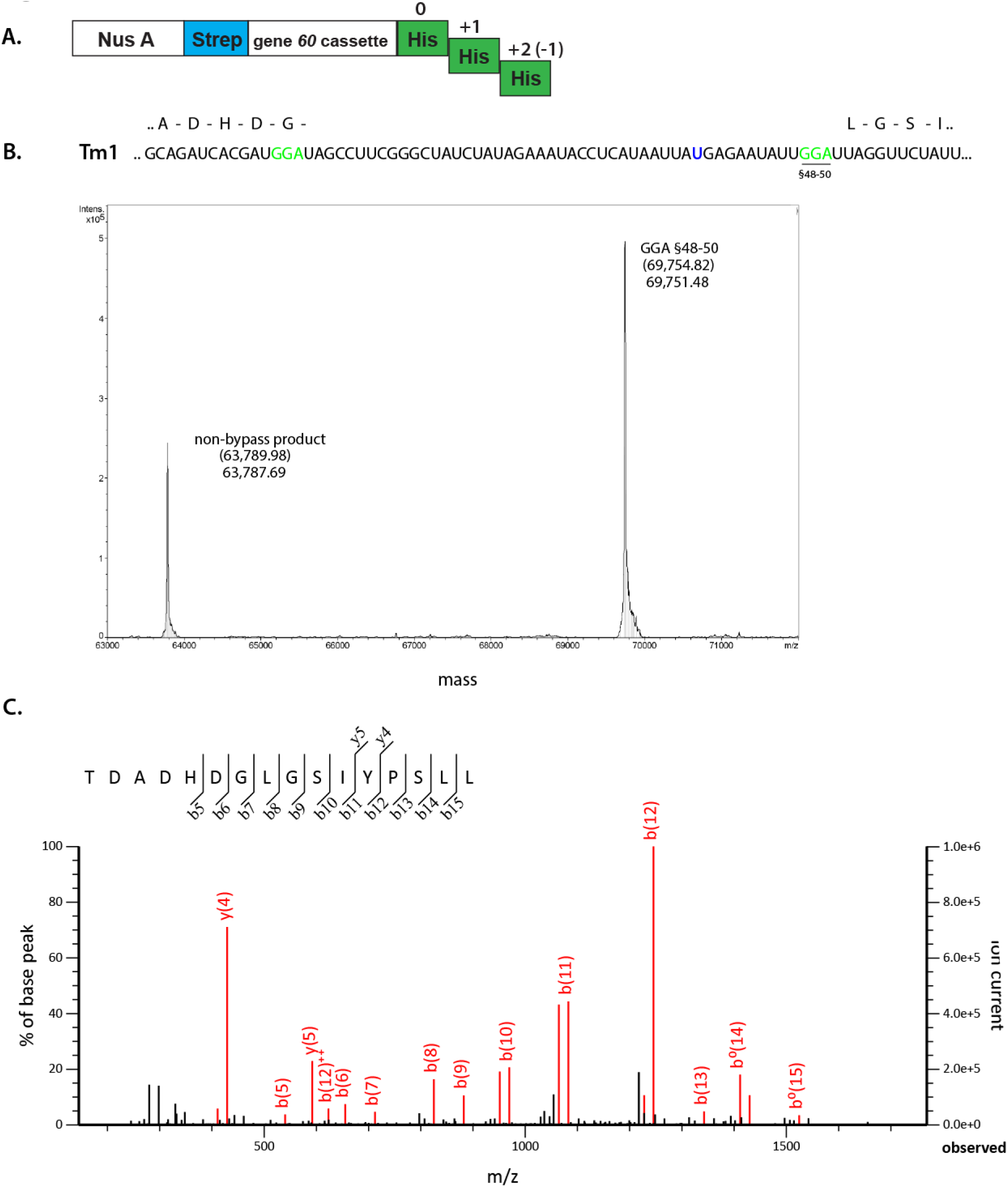
Mass spectrometric analyses of the Tm1 bypass product. **(A)** Schematic **o**f the gene *60* constructs. The Strep tag (in blue) and the 6X His tags (in green) allowed purification of bypass products by affinity chromatography (see methods). **(B)** A region of the mWT Tm1 sequence is shown where the nucleotides at the positions of the WT take-off and landing sites are shown in green. The predicted amino acid sequence of a portion of the bypass product is shown above the RNA sequence. In the electrospray mass chromatogram of the intact product, the mass is shown on the x-axis and the intensity is shown on the y-axis. The bypass product is labelled by its landing position (underlined in the sequence). The non-bypass product is also indicated. The predicted mass of the product is indicated in parentheses above the observed mass. **(C)** MS/MS analysis of the bypass peptide. The landing site and sequence of the bypass peptide is indicated on the graph where the mass to charge ratio (m/z) of each fragmented ion is on the x-axis and the percent of the parent peptide ion (base peak) is shown on the y-axis.

With the WT cassette, detectable landing occurs exclusively at the WT landing site, §48-50 (28). There is a UAA stop codon 36-38 nts in the productive new frame 3’ of the take-off site (i.e. in the −1 frame with respect to initiation). In a first set of constructs this UAA stop codon was eliminated by changing A38 to U (shown in blue in Fig.2 panel B; see Methods) to allow detection of products resulting from landing in the −1 frame within the coding gap. The cassette with this modified WT sequence was named Tm1 (<u>T</u>hree prime, <u>m</u>ass spec), from which the following series of constructs was derived.

Electrospray MS analysis of intact Tm1 product, showed that landing occurred exclusively at the canonical WT landing site GGA §48-50 (Fig. 2 panel B; Table 1; Table S1).

**Table 1.**
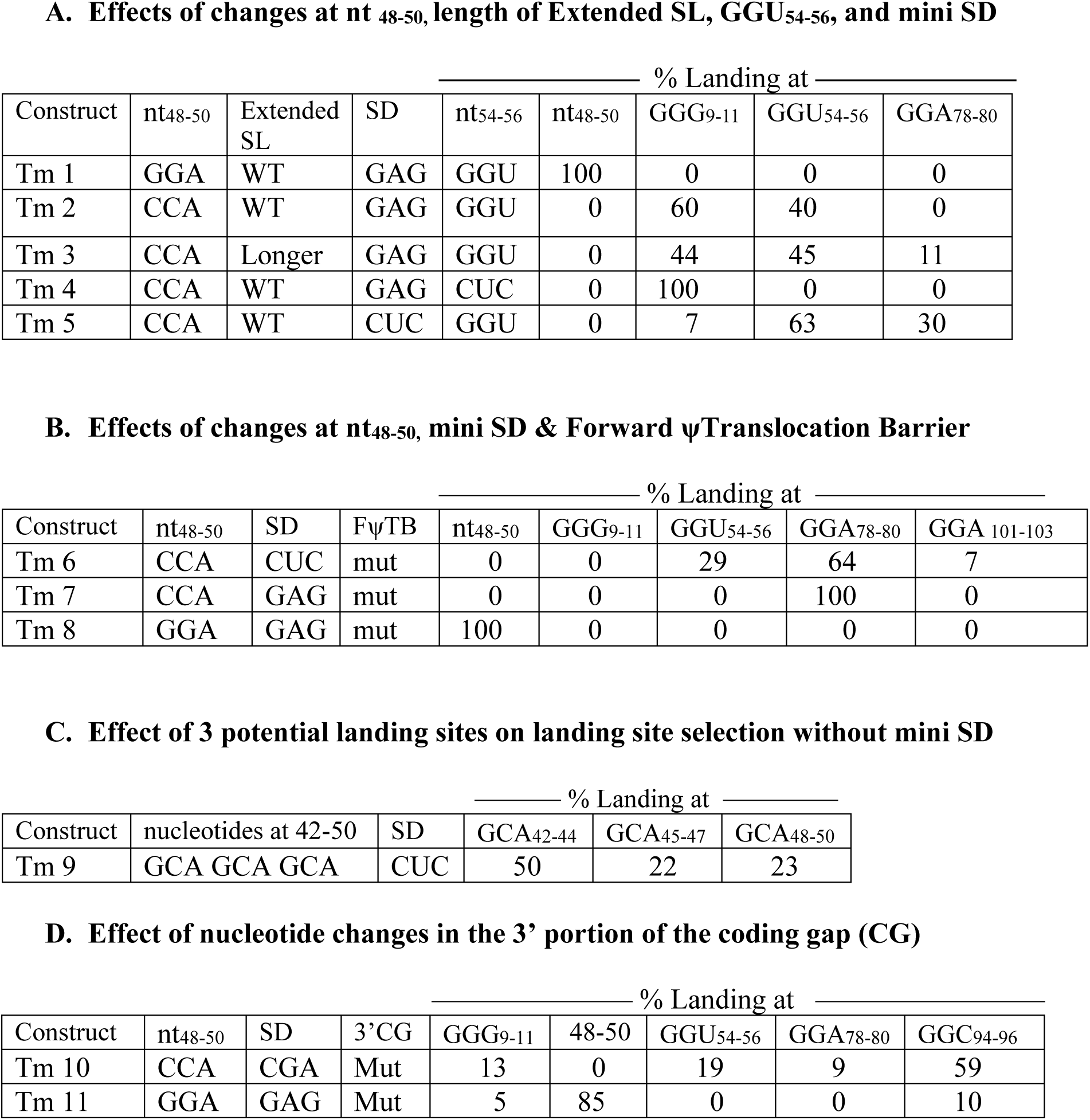
Gene *60* mutant construct features and landing positions utilized.

The result from intact product analysis was confirmed by MS/MS identification of the trans-frame junction peptide (Fig. 2, panel C). Tm1 yielded the same result as authentic WT sequence (28), and the modified WT sequence it contains is referred to as mWT below.

To investigate the effect of eliminating potential re-pairing to mRNA at the canonical WT landing position, construct Tm2 had GGA §48-50 substituted with CCA. Electrospray analysis of intact products shows that 60% of bypass product resulted from landing at GGG §9-11 within the coding gap and 40% at GGU §54-56, 4-6 nt 3’ of the WT landing position (Fig. 3; Table 1; Table S1). MS/MS analysis of chymotryptic peptides confirms that landing occurs at GGG §9-11 and GGU §54-56 (Fig. S1).

**Fig. 3.**
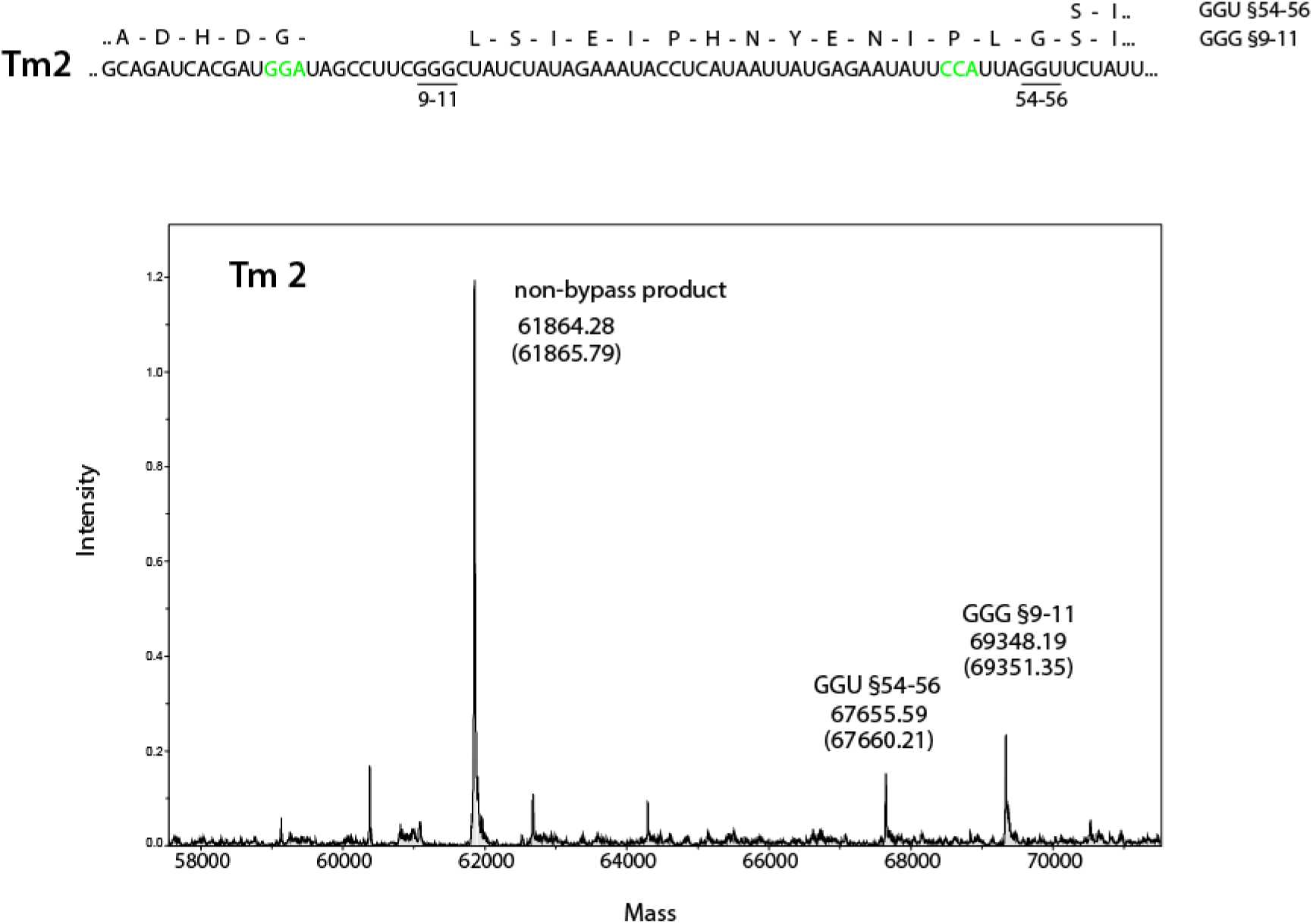
Electrospray mass spectrometric analyses of the Tm2 bypass products. The nucleotides at the “WT” take-off and landing positions are shown in green. The canonical landing position nucleotides were changed to CCA. The predicted amino acid sequences of portions of the observed bypass products are shown above the RNA sequence. In the electrospray mass chromatogram of the intact product, the mass is shown on the x-axis and the intensity on the y-axis. The bypass product is labelled by its landing position (underlined in the sequence). The non-bypass product is also indicated. The predicted masses of the products are indicated in parentheses under the observed mass.

Formation of the Extended SL has been proposed to occur on exit of its component sequence from the mRNA exit tunnel. If formation of the Extended SL provides forward ribosome movement on the mRNA, then a lengthened version may promote bypassing beyond GGU §54-56. Construct Tm3 was designed to test this possibility. Tm3 lengthened the extended SL by 4 base pairs at its base and had the WT landing site replaced by CCA. To elongate the stem, the A, 5 nts 5’ of the take-off codon, was changed to G, and AA, 22 and 23 nts 3’ of the take-off codon, were changed to CG such that there is the potential for 4 additional base pairs at the base of the Extended SL (shown in gray). Mutations are underlined and the WT take-off site is in italics: 5’..ACGCAGATC<u>G</u>CGAT*GGA*TAGCCTTCGGGCTATCTATAG<u>CG</u>ATACCTCATAATT ATGAGAATATTCCA..3’. (In contrast, prior studies with a lengthened SL had the additional base pairs at the top of the stem (33,34, & a proportion in 15); changes at this location would influence formation of both the A-site SL and the Extended SL). Electrospray analysis of intact product showed 44% landing at GGG §9-11, 45% at GGU §54-56 and 11% at GGA §78-80 (Fig. S2 panel A; Table 1; Table S1). MS/MS analysis of chymotryptic peptides confirmed expected junction peptides (Fig. S2 panels B, C, and D). While landing at GGA §78-80 is consistent with a longer extended SL promoting further forward movement, landing at GGG §9-11 is not a simple prediction of this model (see Discussion).

Construct Tm4 (with CCA §48-50 and WT Extended SL) was a derivative of Tm2 but had an alternative landing site utilized in Tm2, GGU §54-56, 3’ substituted with CUC §54-56 (Table 1). The only landing detected by electrospray analysis was at GGG §9-11 (Fig. S3 panel A; Table 1; Table S1). MS/MS confirmed the Electrospray result (Fig. S3 panel B). Removal of an alternative landing site, GGU, 4-6nts 3’ of the WT landing position did not result in landing further 3’.

Construct Tm5 explored the role of the mini-SD sequence GAG §39-41 by having it substituted with CUC. It had CCA §48-50. 30% of the bypass product was due to distant landing at GGA §78-80 (no landing at this site was detected in the presence of the mini-SD). There was increased landing at GGU §54-56 (63%) and reduced (7%) landing at GGG §9-11(Fig. 4 panel A; Table 1; Table S1). Based on earlier work showing a spacing distance-dependent restraining role of an SD sequence (9,35) as discussed below, removal of the restraint was expected to reduce landing at GGG §9-11. [Part of the earlier work referred to explored latitude for programmed frameshifting with ribosomes ultimately moving forwards again. However, only bypassing work can explore disruption of mRNA: rRNA pairing (the extent of the drag) when ribosomes continue to move backwards.]

**Fig. 4.**
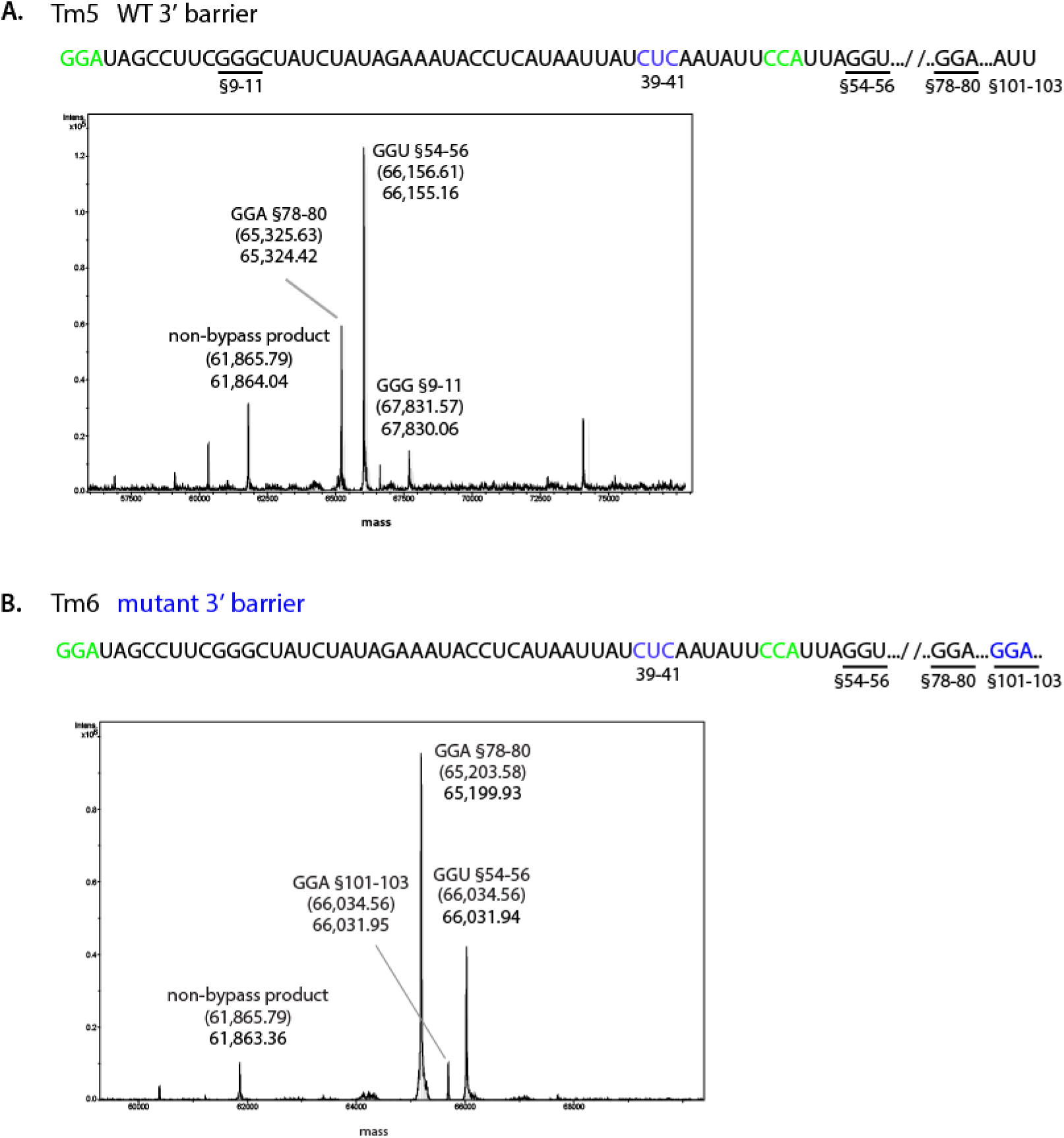
Electrospray mass spectroscopy of intact bypass products. (**A**) A region of the Tm5 is shown where the nucleotides at the positions of WT take-off and landing sites are shown in green and the substitution of the WT SD sequence, GAG 39-41, to CUC is in blue. The sequence of the Forward ψTranslocation Barrier is WT. Bypass products are labeled in the mass chromatogram by their landing positions (underlined in the sequence). The predicted mass for each product indicated in parentheses above the observed mass. The mass of the non-bypass product is also indicated. Unlabeled peaks could not be assigned to any potential bypass product. (**B**) A similar region of Tm6 sequence is shown. It is comparable to Tm2 but contains the mutant forward barrier sequence, X, which contains GGA at positions §101-103 shown in blue. Bypass products are labeled in the mass chromatogram by their landing positions (underlined in the sequence). The predicted mass for each product is indicated in parentheses above the observed mass.

MS/MS data permit confirmatory identification of the junction peptides (Fig. S4). In the absence of a landing site at §48-50 matched to the anticodon of the retained peptidyl-tRNA, lack of restraint by the SD sequence permits longer distance bypassing.

Construct Tm6 was similar to Tm5 in that it also had CCA §48-50, and the mini-SD, GAG §39-41, substituted with CUC. Additionally, it had a mutation, X (see Methods), that disrupts the WT form of the “Forward ψTranslocation Barrier” structure 3’ of the WT landing site (19). The proportion of landing at GGA §78-80 increases from 30% to 64% and 7% derives from landing at GGA §101-103 (GGA §101-103 is part of the mutated sequence X, and is not present in WT) (Fig. 4 panel B; Table 1; Table S1). The proportion at GGU §54-56 decreases from 63% (with Tm5) to 29%, with no detectable landing within the coding gap. MS/MS data confirm identification of the junction peptides (Fig. S5). When the site of landing in WT is not matched to the anticodon of the retained peptidyl-tRNA, greater length bypassing is evident when the Forward ψTranslocation Barrier is mutated. These conditions provide evidence that *in vivo* the WT barrier does indeed impede forward ψtranslocation. With a WT landing site does the barrier facilitate landing at §48-50 at a detectable level? This is addressed with Tm8 below, but we next consider the effect of reintroducing the WT SD.

Construct Tm7 is a modified Tm6 with the WT mini-SD GAG §39-41 restored. It had CCA §48-50 and the same 3’ Forward ψTranslocation Barrier X mutant as Tm6. MS/MS data gives no hint of landing at GGU §54-56, just at GGA §78-80 (Fig. 5; Table 1).

**Fig. 5.**
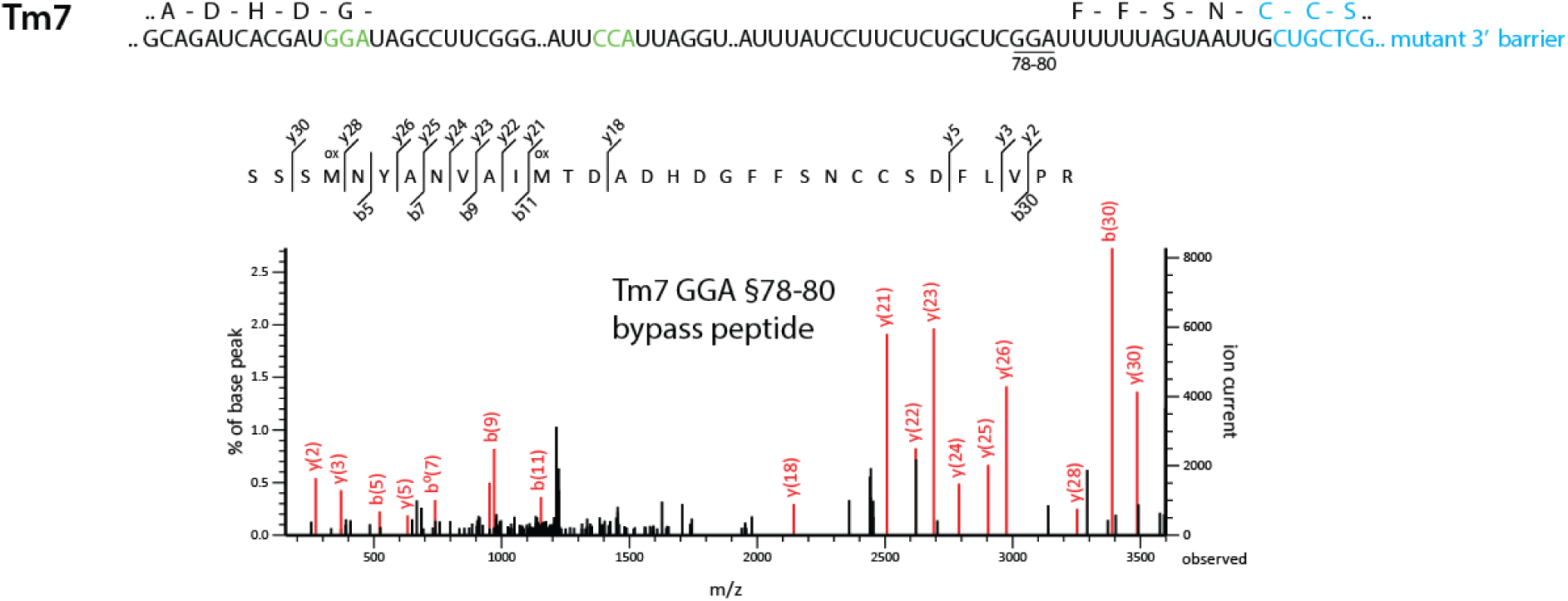
MS/MS analysis of the Tm7 bypass peptide. The RNA sequence of Tm7 shown with the nucleotides at the WT take-off and landing positions in green. The nucleotide substitutions in the 3’ Forward ψTranslocation Barrier are shown in blue. The utilized landing site, GGA 78-80, is underlined and the predicted amino acid sequence of a portion of the bypass product is indicated above the RNA sequence with WT amino acids in black and the amino acids encoded by the mutant RNA sequence in blue. The amino acid sequence of the bypass peptide is indicated in the graph where the mass to charge ratio (m/z) of each fragmented ion is on the x-axis and the percent of the base peak is shown on the y-axis.

Though an SD with GGAGG can still maintain pairing with an anti-SD on movement to 12nts from the P-site and has a restraining effect (9,35,36), the result here with the minimal SD, GAG, indicates no effect of such an SD in directing landing 12 nts 3’. With a GGA take-off site, and with §48-50 being CCA, the Forward ψTranslocation Barrier is deduced to have a greater effect than the WT SD. Consistent with this, the presence of the WT Forward ψTranslocation Barrier with CCA at §48-50 results in no landing further downstream than GGU §54-56 (Fig. S1 panels A and B). Interestingly, landing occurs within the coding gap at GGG §9-11, consistent with backwards ribosome movement due to both the Forward ψTranslocation barrier and an effect of SD: anti-SD pairing dragging back forward moving ribosomes before disruption of the pairing (Fig. S1 panel A).

Construct Tm8 had the WT GGA §48-50 (instead of CCA §48-50), but retained the 3’ forward barrier mutant X. This was only studied by MS/MS chymotryptic peptide analysis. Only a product due to landing at GGA 48-50 was detected (Fig. S6, Table 1). Since Tm8 differs from mWT, Tm1, only by the 3’ forward barrier X mutation, and they yield comparable results, any role for the 3’ structure is subtle.

### Resumption of codon scanning within the coding gap by the anticodon of the retained peptidyl-tRNA

Previously in a mutant with an AUU at the take-off site and 3 tandem AUUs §42-50, landing was only detected at AUU §48-50 (none at either of the two 5’ adjacent AUUs) (28). This led to the inference that with a WT coding gap peptidyl-tRNA anticodon scanning of coding gap sequence did not commence until the site of landing in WT was reached. The alternative explanation, which was not considered in Wills *et al*., (28), is that the SD §39-41 strongly influences landing specifically at the AUU §48-50, 6nts 3’. This contrast prompted the generation of Tm9.

Construct Tm9 has a GCA take-off site and GCAGCAGCA at §42-50 to give stronger codon: anticodon pairing than when the matched take-off and landing codons are AUU (16). In addition, it has the WT SD substituted to CUC §39-41 to preclude an SD influence on landing. Mass spectrometric analysis of Tm9 products yielded several unidentified peaks in the electrospray mass chromatogram, with landing at the GCAs comprising only a very small proportion of the total (Fig. 6; Table S1).

**Fig. 6.**
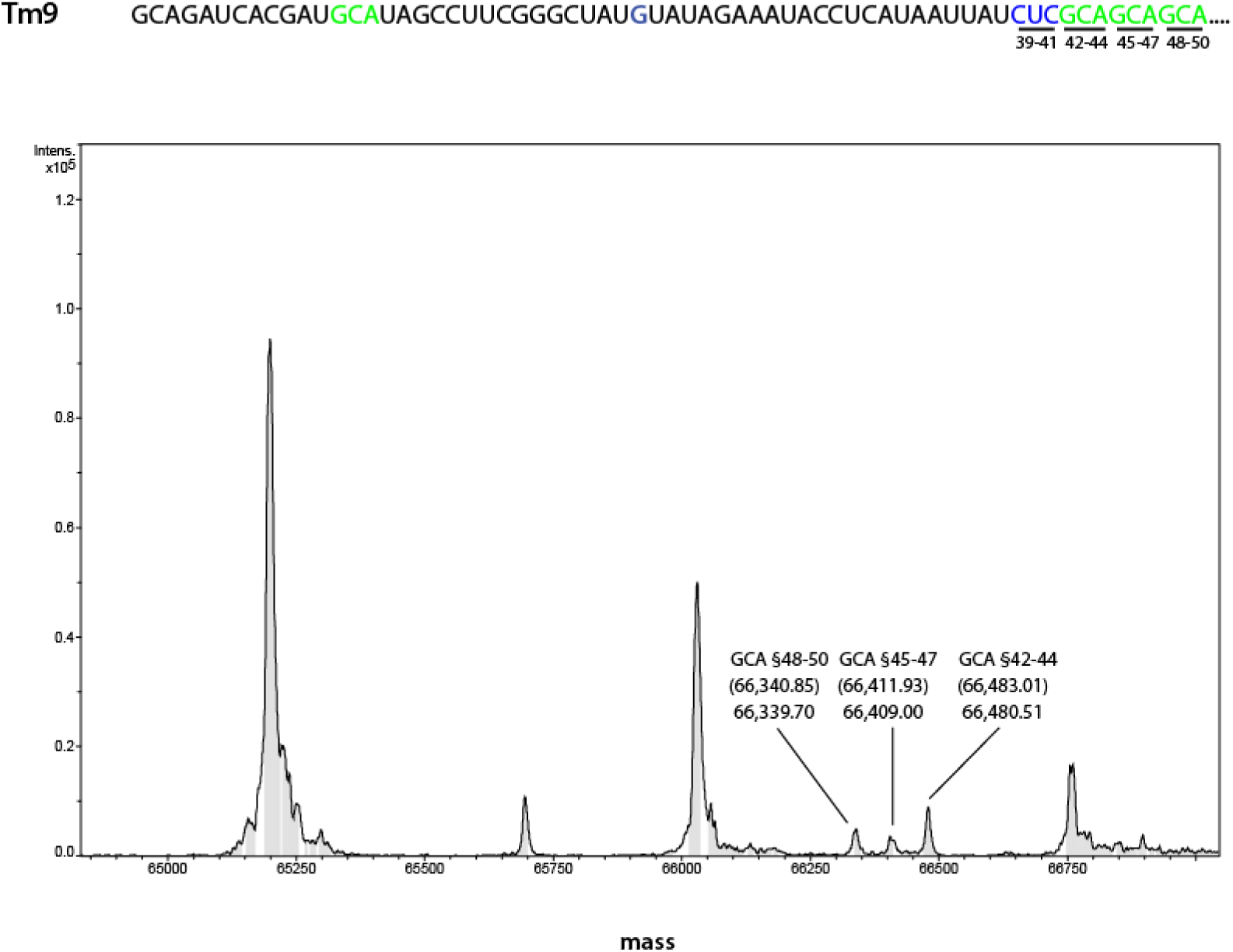
Electrospray mass spectrometric analyses of Tm9 bypass products. A region of the TM9 sequence is shown where the nucleotides, GCA, at the positions of the WT take-off and landing sites are shown in green. In addition, the 6 nucleotides 5’ of the WT landing position were substituted with GCAGCA to create 3 tandem potential landing sites. The nucleotide substitutions introduced to maintain base pairing within the main SL (G) and to eliminate the WT SD sequence (CUC) are shown in blue. In the electrospray mass chromatogram, the mass is shown on the x-axis and the intensity is shown on the y-axis. The bypass products are labelled by their GCA landing positions (underlined in the sequence). The predicted masses of the products are indicated in parentheses above the observed mass. Unlabeled peaks could not be assigned to any potential bypass product.

Of the products that did derive from landing at GCA, 50% were from landing at GCA §42-44, 22% at GCA §45-47 and 23% at GCA §48-50, confirmed by MS/MS analysis (Fig. S7, Table 1). Prior analysis showed that the influence of SD rRNA: mRNA interaction on directing bypassing ribosome resumption of anticodon: codon pairing (landing) 6 nts 3’ was less when the potential anticodon: codon interactions were strong (28). The results presented here that peptidyl-tRNA anticodon scanning starts 5’ of the WT landing site. This brings the results from this approach into line with the earlier conclusions of smFRET studies (21).

### Does nt sequence in the 3’ part of the coding gap other than the SD and landing site affect landing site selection?

Prior work showed that mutants of the take-off stem loop structure can enhance landing at GGG9-11. A search was undertaken for mutants of the 3’ part of the coding gap that enhance landing at other than the WT landing site position (32). One of the mutants, Tm10, is analyzed here. Its sequence is **CGCCAACGACCCGUU** §33-47, CCA §48-50 (substitutions in blue, and WT nts in black). This has no internal SD, but has CCA §48-50 in place of GGA §48-50. The sequence 3’ of §48-50 is WT (Fig. 7; Table 1). It is comparable to Tm5 which has CUC substituted for GAG §39-41 and CCA §48-50, but is otherwise “WT” (Fig. 4, panel A; Table 1).

**Fig. 7.**
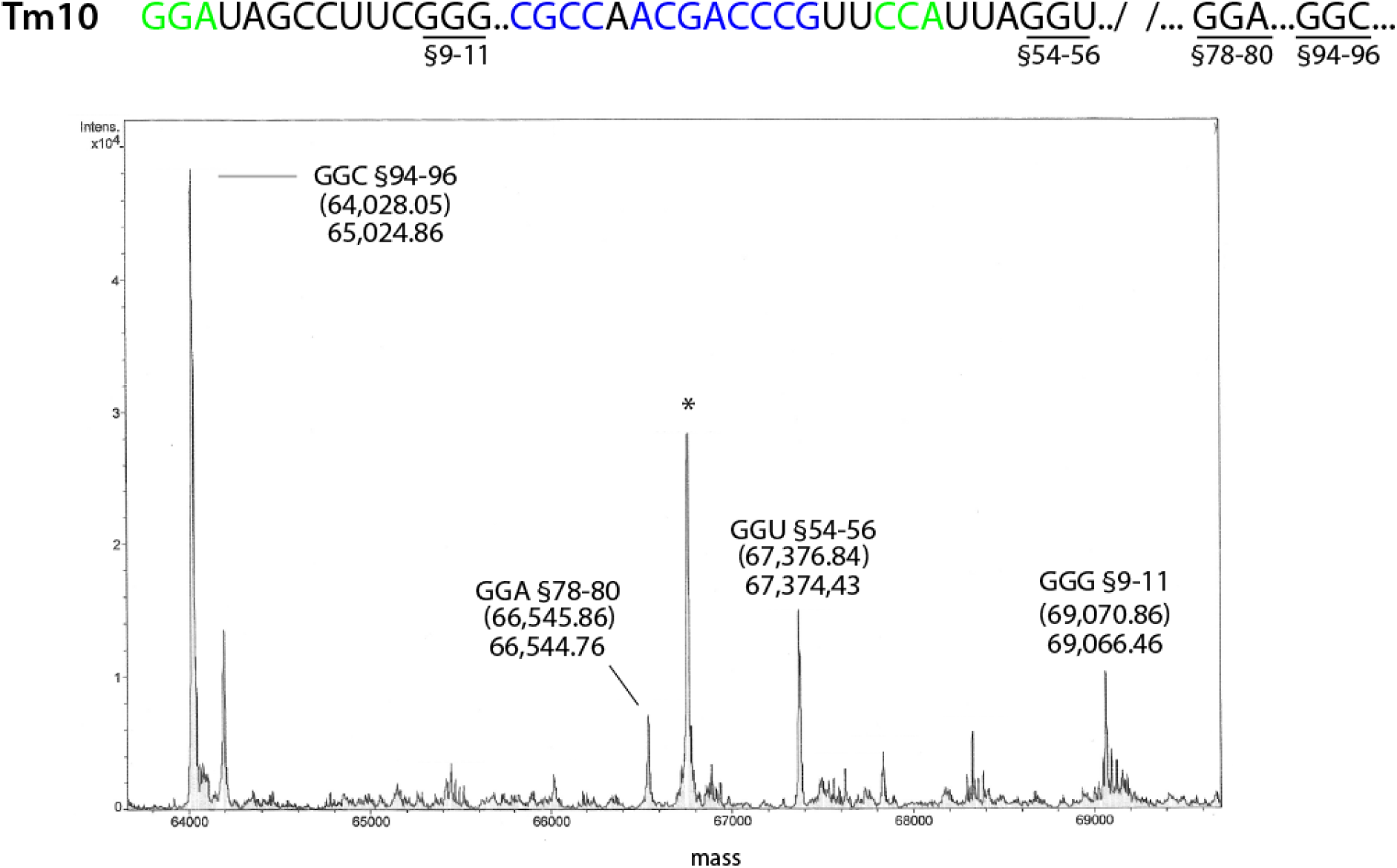
Electrospray mass spectrometric analyses of Tm10 bypass products. A region of the TM10 sequence is shown where the nucleotides at the positions of the WT take-off and landing sites are shown in green. The nucleotide substitutions in the 3’ part of the coding gap are shown in blue. In the electrospray mass chromatogram, the mass is shown on the x-axis and the intensity is shown on the y-axis. The bypass products are labelled by their landing position (underlined in the sequence). The predicted masses of the products are indicated in parentheses above the observed mass. Unlabeled peaks and * could not be assigned to any potential bypass product.

59% of the product from construct Tm10 (Fig. 7; Table 1; Table S1) resulted from landing at GGC §94-96 whereas this product was not detected with Tm5. In Tm 10 landing also occurred at GGU §54-56 (19%), GGA §78-80 (9%) and GGG §9-11 (13%). For Tm5, the corresponding percentages were 63%, 30% and 7% (Fig. 4 panel A; Table 1; Table S1).

Construct Tm11 has the WT landing site, GGA §48-50, as well as having the natural mini-SD, GAG §39-41, reintroduced into the Tm10 context (Fig. 8 panel A; Table 1). This is comparable to Tm1 the mWT sequence whose only bypass product results from landing at GGA §48-50 (Fig. 8 panel B; Fig. 2; Table 1; Table S1).

**Fig. 8.**
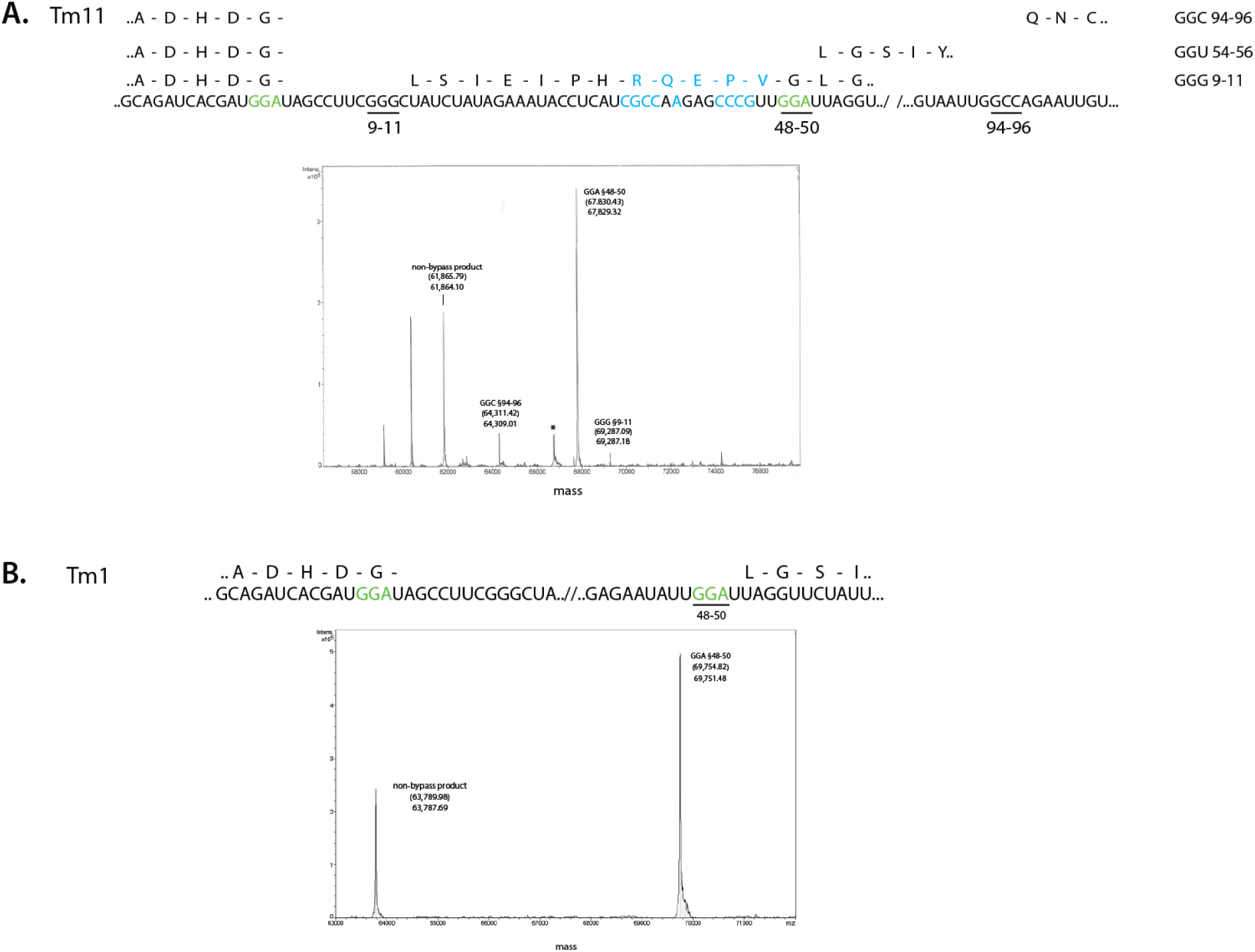
Electrospray mass spectrometric analyses of Tm11 bypass products and comparison with Tm1. **A.** A region of the Tm11 sequence is shown where the nucleotides at the positions of the WT take-off and landing sites are shown in green and the mutant nucleotides in the 3’ part of the coding gap are shown in blue. The predicted amino acid sequences of portions of the bypass products are shown above the RNA sequence with the WT amino acids in black and the amino acids encoded by the mutant RNA sequence in blue. In the chromatogram, the mass is shown on the x-axis and the intensity is shown on the y-axis. The bypass product is labelled by its landing position (underlined in the sequence). The non-bypass product is also indicated. The predicted masses of the products are indicated in parentheses above the observed mass. The peak labelled * (66761 Da) corresponds to the mass of BSA, a component of the cell lysis buffer. Unlabeled peaks could not be assigned to any potential bypass product. **B.** A region of the mWT TM1 sequence is shown where the nucleotides at the positions of the WT take-off and landing sites are shown in green. The predicted amino acid sequence of portion of the bypass product is shown above the RNA sequence. In the chromatogram, the mass is shown on the x-axis and the intensity is shown on the y-axis. The bypass product is labelled by its landing position (underlined in the sequence). The non-bypass product is also indicated. The predicted masses of the products are indicated in parentheses above the observed mass.

With Tm11, the majority of landing (85%) occurs at the same GGA §48-50 (Fig. 8 panel A; Table 1; Table S1). However, landing additionally occurs at GGC §94-96 (10%) and GGG §9-11 (5%) Taken together, Tm10 and 11 show an influence of nucleotide identity in the 3’ part of the coding gap other than the SD and the WT landing site on bypassing distance.

## DISCUSSION

### Forwards bypassing over an extended coding gap: Drop-off

The main part of the work belatedly reported here is to further address a ‘driver’ role for mRNA exiting ribosomes being involved in intra-mRNA pairing. Promotion of unidirectional forward movement of bypassing ribosomes is one consequence. The other is reduced ribosome drop-off while the ‘pushing’ effect is operative. The first pointers to increased drop-off after a ribosome reaches the end of the WT-length coding gap came from experiments where extensive sequence not prone to form secondary structure, was inserted into the coding gap 3’ of nts that form the A-site SL. The efficiency of bypassing dropped disproportionately compared that of the WT coding gap length (18,19). Discovery of the 5’ stem loop (19), led to increasing strong evidence that it caused drop-off in bypassing the coding gap to be lower than what it would be in its absence (19–21). Recent work (8) provides convincing evidence that both the 5’ stem loop and the Extended Stem Loop have this function. (More indirect but supportive evidence from bypassing efficiencies in the present work is not included here.) However, this drop-off lessening function operates only while a bypassing ribosome is traversing the WT length of the coding gap, or actually as recently shown (8) marginally beyond its 3’ end. The proposal coming from several types of analysis, including the present one, is that what is key is the act of ribosome-exiting mRNA becoming involved in intra-mRNA pairing rather than the fully formed structures directly.

How best to construct a bypassing cassette to allow increased intra-mRNA pairing potential, a late-acting ‘Pusher’, that would allow a longer sequence to be bypassed with a low drop-off rate? Bypassing ribosomes in their scanning mode do not cope well with melting coding gap mRNA structure (19), and bioinformatics has revealed prevalence of AU-richness in the main and 3’ part of the coding gap in diverse occurrences of gene *60* bypassing (5). Consequently, it may not be productive to try to create the strong pairing potential fully within the scanned part of the coding gap. However, from what is known it should be innocuous for sequence in the middle of the coding gap that could act as a tetraloop followed by sequence that could act as the 3’ side of a future stem loop. That 3’ side sequence could be designed so that it is complementary to A-site stem loop sequence including GGG_9-11_ (i.e. the 5’ side of a future and new stem loop). The 5’ side of the new stem loop would be unavailable until after the A-site SL sequences has exited the ribosome. Pairing to form the new stem loop should create a new and late acting driver with potential to extend bypassing distance with a low drop-off rate. Construct Tm10, which was obtained in a screen is not ideal for this purpose for several reasons including that it retains the Forward ψTranslocation Barrier. Nevertheless, the results are encouraging for the design of a future construct. Among ribosomes that broke through the 3’ Barrier, more than 6-fold landed at GGC_94-96_ than at GGA_78-80_. In A-site decoding tRNA^Gly^ does not decode GGC (37,38). Ribosomes continuing bypassing to permit landing at GGC_94-96_ (where even in the P-site the third base of GGC is likely not involved in pairing) occurs despite the anticodon of the retained peptidyl-tRNA^Gly^ being matched for GGA_78-80_. Even when the mini-SD is reinserted along with restoration of GGA_48-50_ in its derivative construct Tm11, 10% of the landing still occurred at GGC_94-96_. [Unfortunately, the difficulty of GTP utilization studies limit the practicality of obtaining additional energy utilization data to the extent desirable.] When considering how bypassing is stimulated by recoding signals, it is pertinent to keep in mind that though with much less efficiency, spontaneous long distance bypassing can occur especially under amino acid limitation caused A-site starvation (39,40), a common condition for many bacteria. In contrast programmed gene *60* bypassing does not involve A-site starvation or even A-site access (21,23) due to the locked state of the A-site at the initiation of bypassing (6).

### Unidirectional forward bypassing can transition to a bidirectional mode

The results with Tm1 (WT apart from coding gap UAA_36-38_ being UAU_36-38_) show that landing occurs exclusively at GGA_48-50_. This result was expected from earlier work, though removal of a UAA excludes possible landing that led to termination and lack of detection of a relevant product. This highlights the effectiveness of the combination of recoding signals in ensuring designated landing site fidelity. As several studies have shown, GGG_9-11_ is not scanned by the anticodon of the peptidyl-tRNA retained within the bypassing ribosome while it is en route to the WT landing site, GGA_48-50_.

Our prior mass spectrometric analysis (28) of *in vivo* product from mutants that in the light of subsequent results, have reduced potential for forming a WT-like A-site stem loop, unsurprising led to reduced detectable bypassing and landing at GGG_9-11_, as well as at the canonical WT site. The WT landing site was unaltered in these constructs and also in further mutants with the A-site stem loop unaltered but which disrupt the potential for forming the ‘base region’ (28, Fig 7B) of what is now known as the Extended Stem Loop. This stem loop is proposed to form on mRNA exiting from the ribosome. Bypassing efficiency was also reduced though ~ 70% of landing still occurs at the GGA_48-50_ and ~ 30 % occurs at GGG_9-11_. This is explicable by a model involving formation of the Extended Stem Loop acting as a driving force for bypassing ψtranslocation. On this model the reduced pairing potential of ribosome exiting mRNA would lead to dissipation of the forward driving force 7nts 5’ of its position in WT. The directional freedom results in a proportion of ribosomes moving backwards (5’). Those that do so to the extent that their P-site reaches GGG_9-11_, have to traverse 35nts to do so. Those that continue to move forward 3’ have to do so for just 7nts to reach GGA_48-50_ which as discussed above, has augmentors of landing there. [With a parallel construct with enhanced potential pairing at the base of the extended stem loop, almost all landing was at GGA_48-50_ (28).]

The study here of mutating the site of landing in WT extends this type of analysis. The ribosome cannot know in advance when that landing site is mutated to be non-cognate, as in Tm2. That 60% of the re-pairing to mRNA then occurs at GGG_9-11_, and 40% 3’ of the site of landing in WT, means that after nucleotides 48-50, some of the ribosomes go ‘backwards’ i.e. 5’ as far as nts 9-11 and the rest continue further 3’. This does not occur with WT gene *60*. If a proportion of ribosomes translating the 3’ part of the coding gap of WT gene *60*, was to go backwards as far as GGG_9-11_ with ensuing anticodon: codon pairing, this should have been reflected in the product analyzed. As it is not, we deduce that <u>backwards</u> <u>bypassing</u> on WT gene *60* mRNA *in vivo* <u>does not occur</u> (at a detectable level).

Irrespective of how it occurs, backwards (3’ to 5’) 70S ribosome scanning as far as GGG_9-11_ implies lack of interference by a following ribosome synthesizing the C-terminal region of the nascent peptide signal. We have not tested non-synonymous mutants of the KKYK sequence that at the constriction site of the exit tunnel have a prominent slow-down effect on ribosome movement. Might KKYK interaction with the exit tunnel be relevant to non-occluded mRNA being available for backwards scanning or is its presence incidental? Since such scanning does not occur with the WT T4 gene *60* bypassing cassette, selection for KKYK being present has not been for this reason. However, when the canonical WT landing site is mutated, as a ribosome retraces its steps backwards (5’) from nts 48-50, the Extended SL, which contains GGG_9-11_ is expected to have completely formed outside the mRNA exit site. That peptidyl-tRNA in the backwards moving ribosome can pair with GGG_9-11_ implies that the Extended SL unfolds and mRNA that formed it can re-enter the ribosome enabling GGG_9-11_ to occupy the P-site. If the Extended SL initially serves any backstop function, this function can only be transitory.

In Tm3 the sequence allowed potential base pairing to lengthen the Extended Stem Loop, and also the canonical landing site was replaced by CCA. Unsurprisingly there was substantial landing 3’ of the site of WT landing consistent with the potential for more exiting mRNA forming an elongated Extended stem loop with more prolonged dragging of mRNA out of the ribosome to promote more extensive unidirectional forward bypassing. Nevertheless, it could be regarded as cautionary that 44% of Tm3 productive landing occurred at nts 9-11 (i.e. from ribosomes that moved backwards though the mini-SD present might have had a minor role in this). However, creating the potential of increased Extended Stem Loop pairing may have come at the price of disrupting other relevant pairing. Rather than independent formation of two discrete stem loops, the 5’SL (or Initiating Driver SL) and the Extended SL (or Main Driver SL), it seems that uninterrupted pairing of mRNA exiting the ribosome would seem more favorable for a forward ‘driver’ mechanism. While the potential for such continuous pairing for T4 gene *60* is not obvious, we propose that it occurs and it will be addressed in future work. If correct, then mutants that extend the base of the Extended stem loop may interfere with potential pairing that occurs in the transition from the 5’ stem loop to the Extended stem loop. Indeed, NMR and other approaches have shown that the 5’ stem loop especially its ‘lower’ part, is not stable (18,21) consistent with lack of stability freeing constituent segments for pairing with further 3’ mRNA as it in turn emerges from the ribosome. The selective advantage for pairing of exiting mRNA would only be expected to be sufficient to push the ribosome until the ribosomal P-site reaches the landing site, but not beyond that. Dissipation of the driving force could then give bypassing directional latitude. [In addition to issues related to the formation of a sequential series of multiple, relatively small structures influencing bypassing, there are likely limits to the extent of the potential for which pairing involving a single extensive stem loop structure could ‘drive’ unidirectional forward bypassing over much longer distances. Considerations include potential susceptibility to double-stand specific nucleases and effects of preformed structures ahead of an approaching ribosome. Regarding the latter, as introduced above, effects of even the WT mRNA structures have become evident (8).] Because of the inter-relatedness of several of the bypassing recoding signals, inferences only from mutants need to be treated cautiously. However, as the prime focus of the present work is on substitution of the site of landing in WT and not on modification of important RNA structures, and as it is being complemented by emerging kinetic analyses, such concerns are minimized.

Previously a mutant gene *60* cassette was generated to address whether bypassing involving decoding resumption 5’ of the take-off site and in a different frame could be detected. Would a translating ribosome read a short sequence first in one frame and then go backwards to read the same sequence in an alternative frame to synthesize a single polypeptide? However, though such backwards bypassing was detected (28), that study did not address the driving force for WT gene *60* unidirectional forward bypassing.

An expectation of the forward-pushing model is that its effect ends as the peptidyl-tRNA anticodon pairs with the landing site, nts 48-50. At that point 5’ of nt 48 there are 26 nts to the predicted 3’ base of the Extended stem loop. Given uncertainty about the number of mRNA nucleotides expected to be between the P-site and exiting mRNA within a bypassing ribosome, this may appear to create a dilemma for a model involving pairing of exiting mRNA influencing directionality to the end of the coding gap. This prompts comparison to the number of nucleotides 5’ from the take-off site to the 5’ side of the tetraloop sequence of the 5’ stem loop – it is also 26. The specific tetraloop sequence present confers significant stability (41) and presumably needs to be outside the ribosome to nucleate formation of the 5’ SL. The start of formation of that 5’ stem loop has been proposed to be important for the initial ψtranslocation steps (7,21) with the degree of its influence being at least partly revealed by the extent to which monosome-mediated bypassing is higher than that of polysome-mediated bypassing (20). In a ribosome profiling experiment employing micrococcal nuclease, mRNA fragments emanating from when the take-off codon was “in” the ribosomal P-site, had two different 5’ ends. The more abundant class has 24-nt and the less abundant class has 40nt, 5′ of the coding gap (20). How the hyper-rotated state of the ribosome may influence cleavage of paired mRNA is difficult to predict. However, given the extent of the circumstantial evidence for pairing of exiting mRNA serving a ribosome forward driver function, and the calculation above of the apparent number of nucleotides between the 3’ end of such structure and the landing site, arguably the main unsolved problem in gene *60* bypassing is the nature of events near the site of mRNA exit.

### Augmentors and inhibitors of anticodon: codon pairing for (WT) landing site selection

The results with Tm5 to Tm8 show that both the mini SD 6’ nts 5’ of nts 48-50 and the Forward ψTranslocation Barrier 3’ of nts 48-50 play a role in directing landing to the WT landing site. While this is evident when nts 48-50 are not cognate for the peptidyl-tRNA anticodon, it is likely they also have subtle effects with the WT landing site, GGA, that has strong re-pairing potential. This conclusion is supported by previous work where the take-off codon was changed to either UCC or GAG and landing at respective matched triplets, natural or created within sequence that in WT would be the coding gap (28), even though the spacing 3’ to what we now know as the Forward ψTranslocation Barrier /3’SL was much greater than in WT. These results contrast with the result from the *in vitro* system in which the Forward ψTranslocation Barrier (FTB) was first identified and found to have a major effect (19). This is interesting though caution is needed because of the different constructs utilized. It seems unlikely the explanation stems from stalling of a further forward ribosome under polysome conditions *in vivo* that were not present with the much lower ribosome loading in the *in vitro* system used. Future experiments to discern if mRNA binding protein(s) or a small RNA not present in the *in vitro* system may be relevant to the contrast. Nevertheless, though the extent of the effect of the Forward ψTranslocation Barrier in the work being reported here is modest, it is greater than that of the mini-SD which exerts a dragging effect on ribosomes as their P-site arrive at the Landing site.

However, the effect of a mini-SD of the same sequence and spacing is much greater when a landing site that only allows significantly weaker anticodon re-pairing to mRNA. Even when there is no potential W-C or wobble pairing coding resumption, though at a low level of coding resumption can still occur (31). [The mini-SD functions as the counterpart of the Arrestor gear on an aircraft carrier for a plane landing on it.] In the WT coding gap, the nucleotides at 42 to 47 are AUUAUU. Consequently, a gene *60* cassette which has substitutions so that the take-off codon is AUU and an AUU at the position_48-50_ allows the choice of three adjacent landing sites. This construct showed landing only at the third AUU, that at 48-50 (28). (A construct with an AAA take-off codon and a short run of As in the vicinity of the landing site, revealed some single nucleotide latitude.) Though we initially interpreted these results incorrectly (28), they are due to the mini-SD 6nt 5’ of it having a strong effect on the weak landing site AUU at that specific 6nt distance. [The initial smFRET study (21) showed that the 3’ part of the coding is scanned.] Further to the discovery of anti-SD sequences in translating ribosomes scanning mRNA for potential complementarity, such interactions may have been relevant for mRNA retention in primordial protein synthesis (42).

### Closing Perspective

Within the past decade, application of smFRET and other relatively new kinetics advances have added enormously to our understanding of gene *60* bypassing. Despite the ability of cryoEM to currently reveal key features, its resolution of events at the site of mRNA exit is limited. As we have attempted to demonstrate here, re-application of mass spectrometric analysis of products synthesized by mutants of bypassing components can still sensitively play a role in the combined approaches needed for a fuller understanding. However, input from other techniques perhaps including chemical probing combined with next generation sequencing (43,44) are needed. [The only reported probing experiments for T4 gene *60* mRNA were informative, though they were performed in the absence of ribosomes (45).]

The results here prompt consideration of the term ‘phaseless wandering’ coined in 1967 (46) in studies of the ‘rapid lysis’, rII gene (47) that is almost adjacent to gene *60* in which the bypassing we are studying occurs. [Their proposal for termination and linked reinitiation involving at least one subunit that did not dissociate from mRNA was later strongly supported (48). However, for one mutant (involving terminator 360_UGA_ which involves a new AUG in the sequence C <u>GAA</u> UGA GCU <u>GAA</u>) rather than involving reinitiation at a wrong frame AUG we consider it more likely that a short ‘hop’ from the GAA 5’ adjacent to a stop codon was involved.] Though great genetics was involved in the work that led to the term ‘phaseless wandering’, there is a big contrast to the scanning by whole 70S ribosomes in ψtranslocation mode in gene *60* bypassing (7). The latter involves single nucleotide shifts to give anticodon sampling of each overlapping triplet with utilization of much more GTP than standard translation. It is far from passive sliding. Relevance of bypassing findings to wider aspects of gene expression merit future discussion.

## ACKNOWLEDGEMENTS

Highly appreciated support from Ray Gesteland made this work, which was completed several years ago, possible. We are grateful to Jody Puglisi’s insight about the relevance of landing site codon pairing strength. We thank Gerard Manning, Michael O’Connor, Ahmad Jomaa and Robert (Bob) Weiss for their comments on the ms, to Ekaterina (Katya) Samatova for her thoughts about the contrasting 3’SL results and Kate O’Connor for modifying a figure. We appreciate Michelle O’Connor’s role in precursor work to isolate the 3’ gap mutant and Sinéad O’Loughlin’s related work. J.F.A. was supported by Science Foundation Ireland grant 13/1A/1853 and during ms write-up by a following grant, Irish Research Council Advanced Laureate award IRCLA/2019/74.

## AUTHOR CONTRIBUTIONS

NMW and J.F.A. planned the work. NMW performed all the experiments. KP performed the mass spectrometry and helped in its interpretation. JFA wrote the ms with input from NMW.

