## Supplementary material for "Evidence for the pairing of mRNA exiting ribosomes acting as a Driver of unidirectional forward bypassing: Its cessation permits Backwards scanning"

### SUPPLEMENTARY Table and Figures

#### Table S1

| Construct | Landing site | % Bypass product | Obs. MW | Pred. MW | % mass difference |
|---|---|---|---|---|---|
| **Tm1 (“WT”)** | | | | | |
| | GGA §48-50 | 100 | 69,751.48 | 69,754.80 | 0.005 |
| **Tm2 (CCA §48-50)** | | | | | |
| | GGG §9-11 | 60 | 69,348.19 | 69,351.00 | 0.004 |
| | GGU §54-56 | 40 | 67,655.59 | 67,660.21 | 0.007 |
| **Tm3 (extended main SL)** | | | | | |
| | GGU §54-56 | 45 | 66,172.27 | 66,175.66 | 0.006 |
| | GGG §9-11 | 44 | 67,804.40 | 67,808.54 | 0.005 |
| | GGA §78-80 | 11 | 65,339.14 | 65,334.68 | 0.007 |
| **Tm4 (CCA §48-50, CUC §54-56)** | | | | | |
| | GGG §9-11 | 100 | 69,402.90 | 69,407.46 | 0.007 |
| **Tm5 (CCA §48-50, CUC §39-41)** | | | | | |
| | GGU §54-56 | 63 | 66,155.16 | 66,156.61 | 0.002 |
| | GGA §78-80 | 30 | 65,324.41 | 65,325.63 | 0.002 |
| | GGG §9-11 | 7 | 67,830.06 | 67,831.57 | 0.002 |
| **Tm6 (CUC §39-41, X seq 3’)** | | | | | |
| | GGA §78-80 | 64 | 65,199.93 | 65,203.58 | 0.006 |
| | GGU §54-56 | 29 | 66,031.95 | 66,034.56 | 0.004 |
| | GGA §101-103 | 7 | 65,696.00 | 65,699.86 | 0.006 |
| **Tm7 (CCA §48-50, X seq 3’)** | | | | | |
| | GGA §78-80 | | NA | | |
| **Tm8 (GGA §48-50, X seq 3’)** | | | | | |
| | GGA §48-50 | | NA | | |
| **Tm9 (GCA take-off, GCAGCAGCA §42-50)** | | | | | |
| | GCA §42-44 | 50 | 66,480.51 | 66,483.01 | 0.004 |
| | GCA §48-50 | 23 | 66,339.70 | 66,340.85 | 0.004 |
| | GCA §45-47 | 22 | 66,409.00 | 66,411.93 | 0.002 |
| **Tm10 (CCA §48-50, mutant 3’ gap seq)** | | | | | |
| | GGC §94-96 | 59 | 64,024.86 | 64,028.05 | 0.005 |
| | GGU §54-56 | 19 | 67,374.43 | 67,376.84 | 0.004 |
| | GGG §9-11 | 13 | 69,066.46 | 69,070.86 | 0.006 |
| | GGA §78-80 | 9 | 66,544.76 | 66,545.86 | 0.002 |
| **Tm11 (GGA §48-50, GAG §39-41, mutant 3’ gap seq)** | | | | | |
| | GGA §48-50 | 85 | 67,829.32 | 67,830.43 | 0.002 |
| | GGC §94-96 | 10 | 64,309.01 | 64,311.42 | 0.004 |
| | GGG §9-11 | 5 | 69,287.18 | 69,287.09 | <0.001 |

Table S1: Compilation of ESI Mass Spectrometry data. Abbreviations: Obs, observed; Pred., predicted; NA, not available (computer issue); % mass difference between predicted and observed masses.

Figure S1

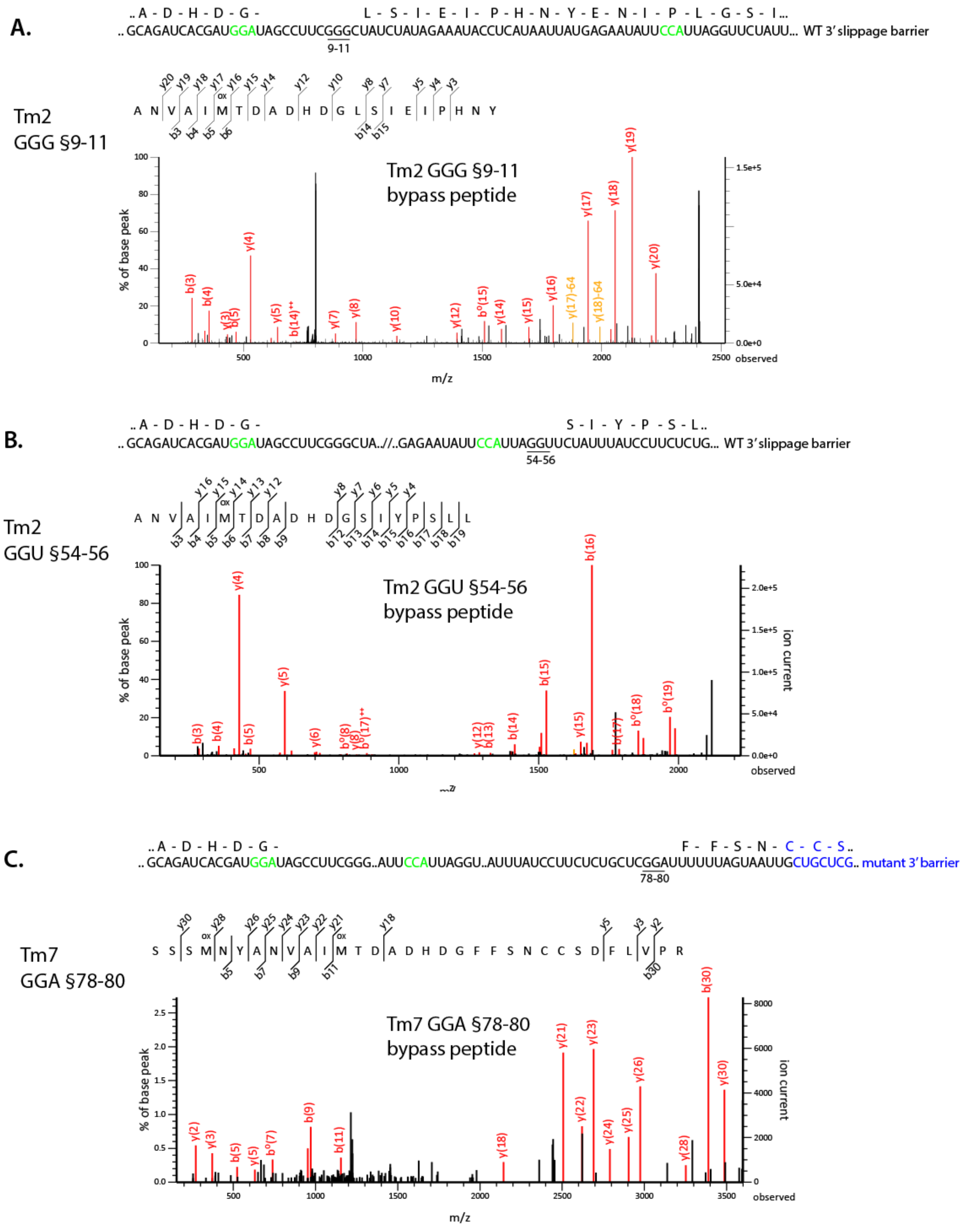


**Fig. S1. MS/MS analysis of bypass (junction) peptides for Tm2 and Tm7.** (**A**) A region of the Tm2 sequence is shown. The nucleotides at the positions of WT take-off and landing sites are shown in green and the sequence of the Forward ψTranslocation Barrier is WT. The underlined sequences are the positions of landing deduced by MS/MS analysis of the bypass products, GGG §9-11 (**A**) and GGU §45-56 (**B**) The mass to charge ratio (m/z) is on the x-axis for each fragmented ion and the percent of the parent peptide ion (base peak) is shown on the y-axis. The values are compiled in a chart in Supplemental Figure 10. (**C**) A similar region of Tm7 sequence is shown. It is comparable to Tm2 but contains a mutant, X, of the Forward ψTranslocation Barrier sequence designed to disrupt its structure. The bypass product deduced by MS/MS analysis results from landing at GGA §78-80, overlined. The mass to charge ratio (m/z) is on the x-axis and the percent of the base peak is shown on the y-axis.

Figure S2

**Tm3** ...AUCGCGAUGGAUAGCCUUCGGGCUAUCUAUAGCGAUACCUCAUAAUUAUGAGAAUAUUCCAUUAGGU.../ /...GGA...

§9-11 §54-56 §78-80

**A.**

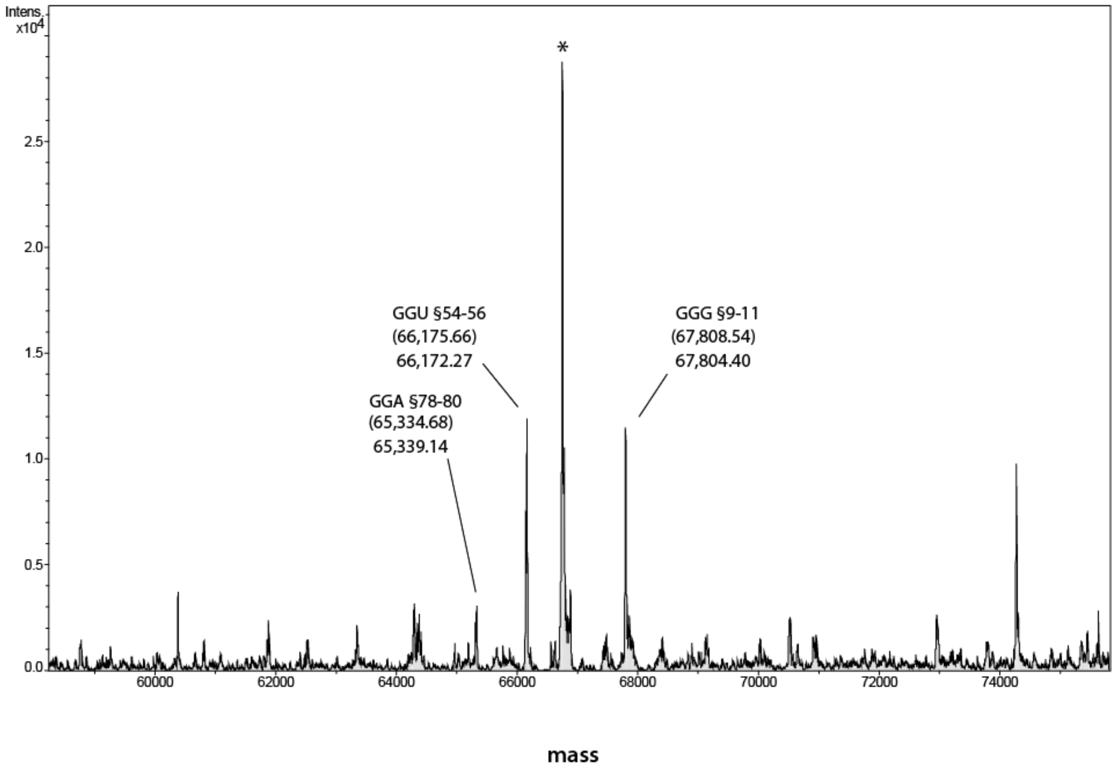

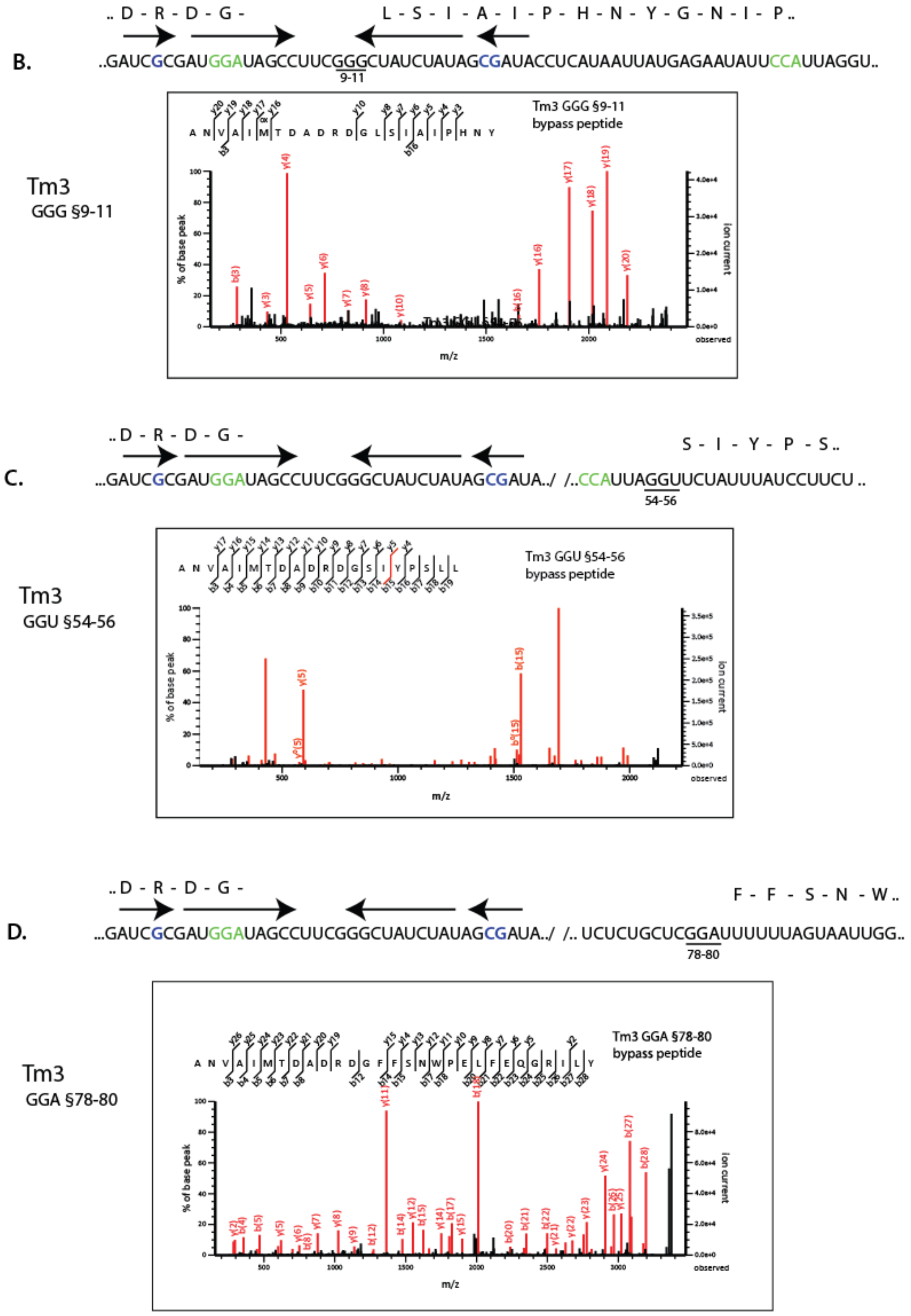


**Fig. S2. Mass spectrometric analyses of Tm3 bypass products. (A)** A region of the Tm3 sequence is shown where the nucleotides at the positions of the WT take-off and landing sites are shown in green. The nucleotide substitutions introduced to extend base pairing at the base of the main SL are shown in blue and the regions predicted to pair are overlined with arrows. In the electrospray mass chromatogram, the mass is shown on the x-axis and the intensity is shown on the y-axis. The bypass products are labelled by their landing position (underlined in the sequence). The predicted masses of the products are indicated in parentheses above the observed mass. The peak labelled * (66761 Da) corresponds to the mass of BSA, a component of the cell lysis buffer. Unlabeled peaks could not be assigned to any potential bypass product. **(B, C and D)** MS/MS analyses of the three bypass peptides showing the sequence of each peptide and the landing site utilized, §GGG 9-11, GGU §54-56 and GGA §78-80 respectively. In the graphs, the mass to charge ratio (m/z) of each fragmented ion is on the x-axis and the percent of the base peak is shown on the y-axis.

Figure S3

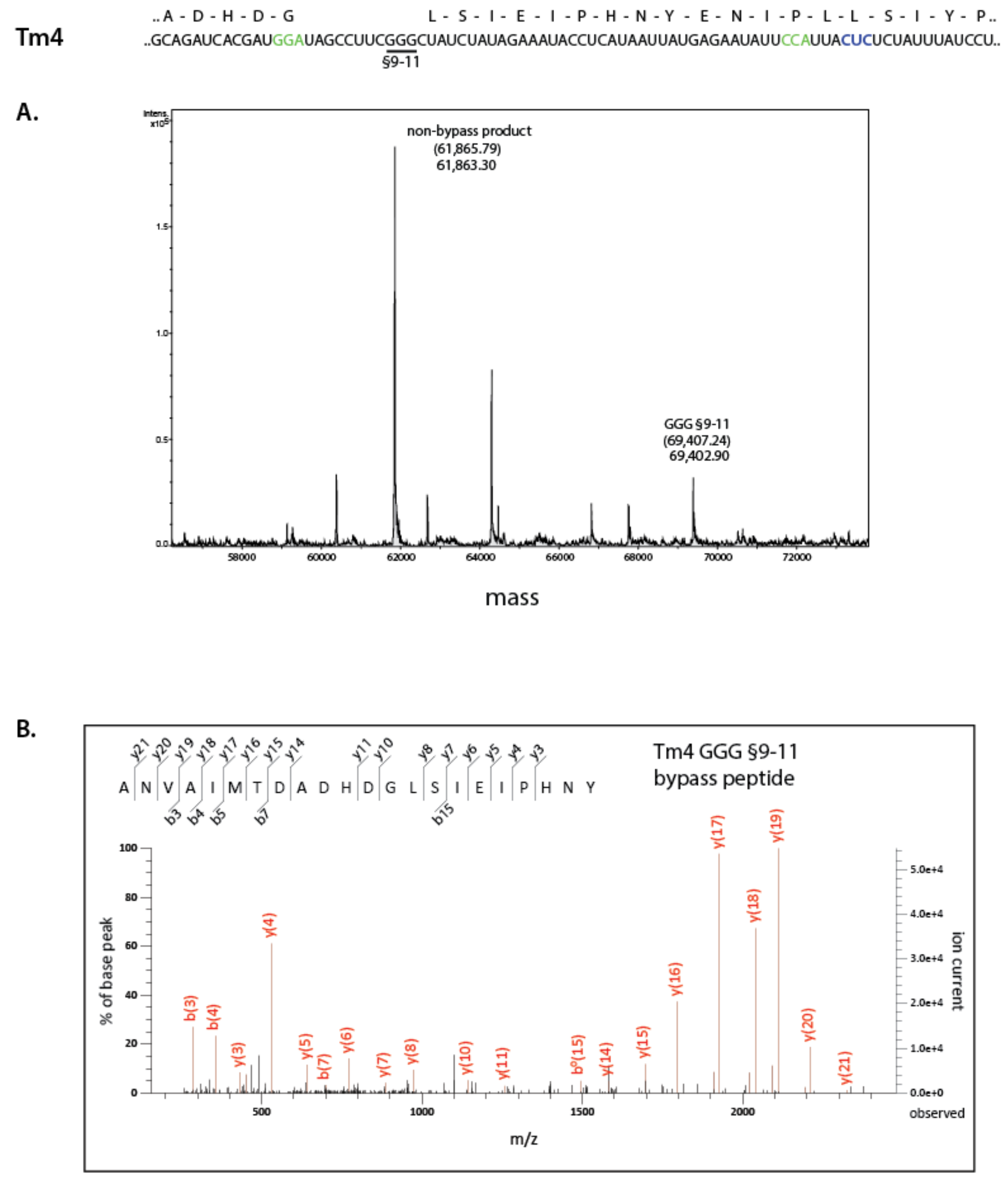


**Fig. S3. Mass spectrometric analyses of the Tm4 bypass product**. **(A)** A region of the Tm4 sequence is shown where the nucleotides at the positions of the WT take-off and landing sites are shown in green. It is similar to Tm2, but has CUC substituted at positions §54-56 (GGU in Tm2 that was utilized as a landing site) shown in blue. In the electrospray mass chromatogram of the intact product, the mass is shown on the x-axis and the intensity is shown on the y-axis. The bypass product is labelled by its landing position (underlined in the sequence). The non-bypass product is also indicated. The predicted masses of the products are indicated in parentheses above the observed mass. Unlabeled peaks could not be assigned to any potential bypass product. **(B)** MS/MS analysis of the bypass peptide. In the graph, the mass to charge ratio (m/z) of each fragmented ion is on the x-axis and the percent of the base peak is shown on the y-axis.

Figure S4

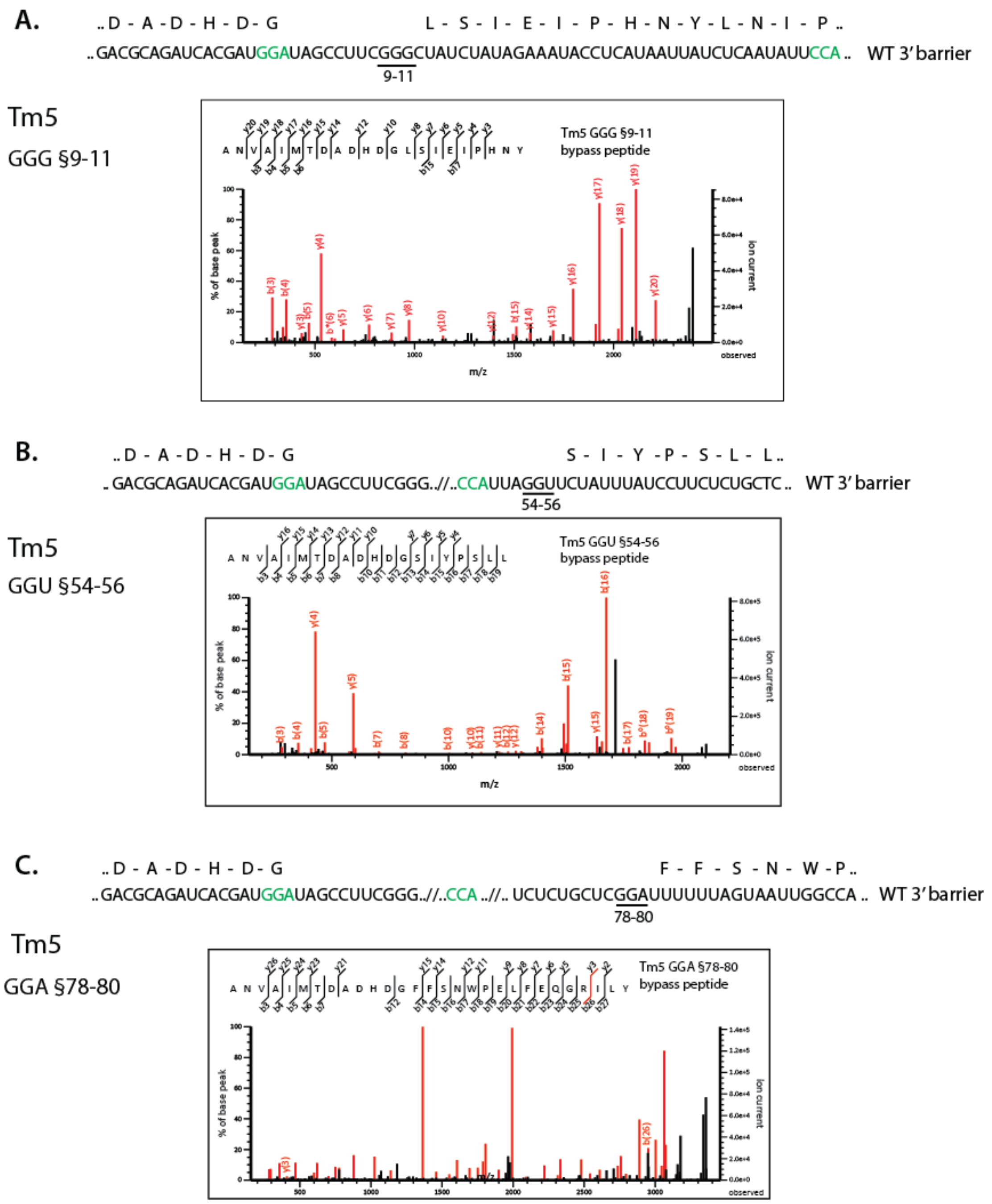


**Fig. S.4 MS/MS analysis of Tm5 bypass products**. **(A)** MS/MS analysis of the GGG §9-11 bypass peptide. In the graph, the mass to charge ratio (m/z) of each fragmented ion is on the x-axis and the percent of the base peak is shown on the y-axis. **(B)** MS/MS analysis of the GGU §54-56 bypass peptide. In the graph, the mass to charge ratio (m/z) of each fragmented ion is on the x-axis and the percent of the base peak is shown on the y-axis. **(C)** MS/MS analysis of the GGA §78-80 bypass peptide. In the graph, the mass to charge ratio (m/z) of each fragmented ion is on the x-axis and the percent of the base peak is shown on the y-axis. (The relevant RNA sequence of Tm5 and corresponding electrospray mass chromatogram are shown in Fig. 4, panel A.)

Figure S5

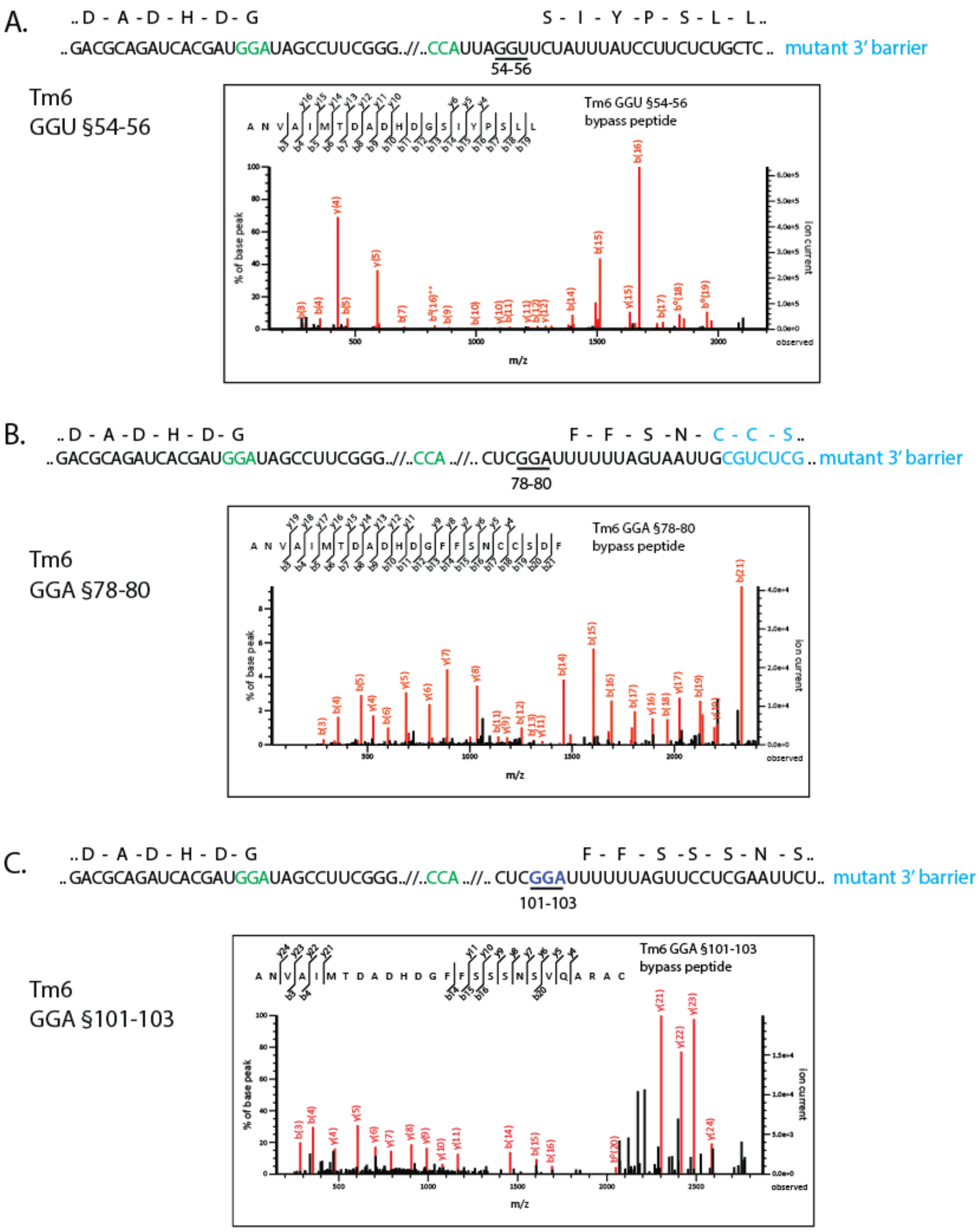


**Fig. S.5 MS/MS analysis of Tm6 bypass products**. **(A)** A portion of the Tm 6 sequence is shown with the WT take-off and landing positions in green. The mutant 3' Forward ψTranslocation Barrier is indicated in blue. The predicted amino acid of the §GGU54-56 bypass product is shown above the RNA sequence. The landing site is underlined. MS/MS analysis of the GGU §54-56 bypass peptide. In the graph, the mass to charge ratio (m/z) of each fragmented ion is on the x-axis and the percent of the base peak is shown on the y-axis. **(B)** A portion of the Tm 6 sequence is shown with the WT take-off and landing positions in green. The predicted amino acid of the §GGA 78-80 bypass product is shown above the RNA sequence. The amino acids encoded by the mutant 3' barrier sequence are shown in blue. The landing site is underlined. MS/MS analysis of the GGA §78-80 bypass peptide. In the graph, the mass to charge ratio (m/z) of each fragmented ion is on the x-axis and the percent of the base peak is shown on the y-axis. **(C)** A portion of the Tm 6 sequence is shown with the WT take-off and landing positions in green. The predicted amino acid of the §GGA 101-103 bypass product is shown above the RNA sequence. The §GGA 101-103 landing site, underlined, is part of the mutant 3' Forward ψTranslocation Barrier sequence and is shown in blue. MS/MS analysis of the GGA §101-103 bypass peptide. In the graph, the mass to charge ratio (m/z) of each fragmented ion is on the x-axis and the percent of the base peak is shown on the y-axis. (The sequence of Tm6 and corresponding electrospray mass chromatogram is shown in Fig. 4, panel B.)

Figure S6

Tm8

..A - D - H - D - G - L - G - S - I - Y..

.. GCAGAUCACGAUGGAUAGCCUUCGGGCUAUCUAUAGAAAUACCUCAUAAUUAUGAGAAUAUUGGAUUAGGUUCUAUUUAU... mutant 3' barrier

39-41 §48-50

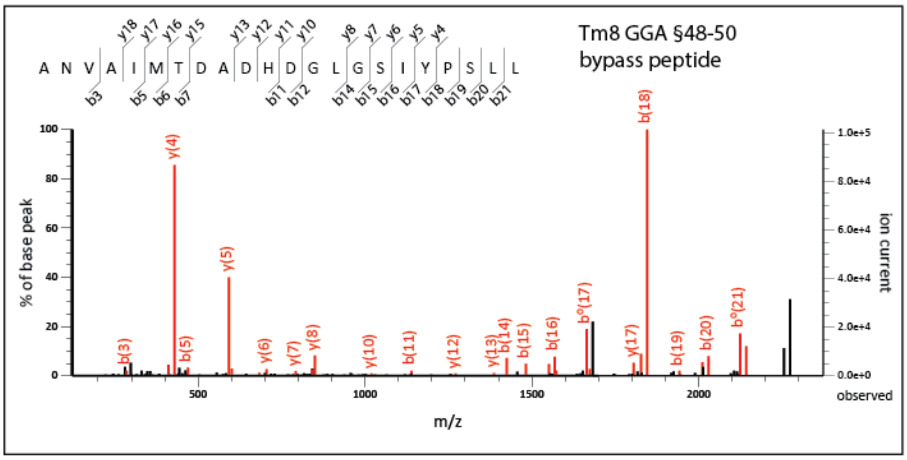


**Fig. S.6 MS/MS analysis of the Tm8 bypass peptide.** RNA sequence of a portion of the Tm 8 sequence is shown. The nucleotides at the WT take-off and landing sites are shown in green. The WT mini SD is present at positions 39-41. The predicted amino acid of a region of the product of landing at GGA §48-50 is indicated above the RNA sequence. MS/MS analysis of the GGA §48-50 bypass peptide. In the graph, the mass to charge ratio (m/z) of each fragmented ion is on the x-axis and the percent of the base peak is shown on the y-axis.

Figure S7

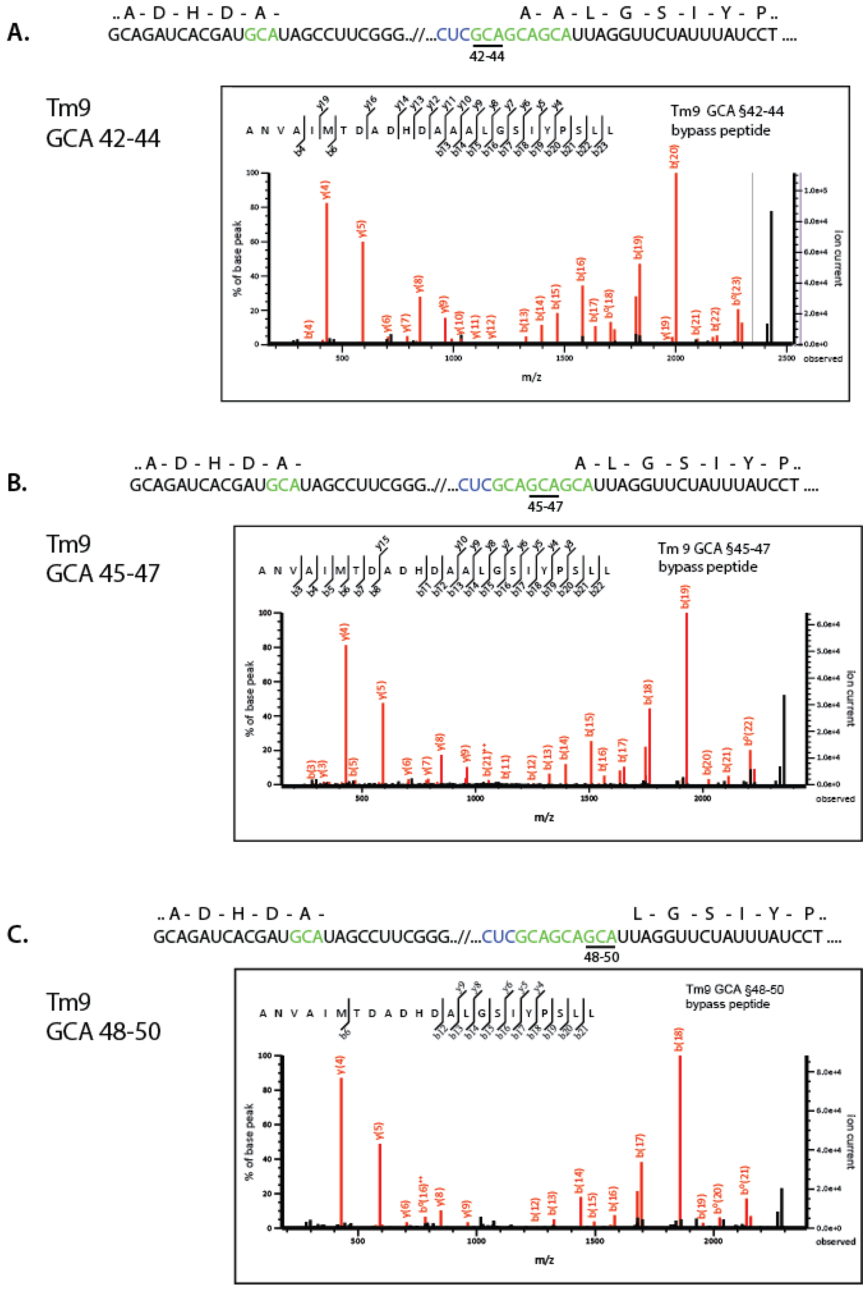


**Fig. S7 MS/MS analysis of the Tm9 bypass peptides. (A)** RNA sequence of a portion of the Tm 9 sequence is shown. The nucleotides at the WT take-off and 3 potential tandem landing sites are shown in green. The WT mini SD was substituted with CUC 39-41 in blue. The predicted amino acid of a region of the product of landing at GCA §42-44 is indicated above the RNA sequence. MS/MS analysis of the GCA §42-44 bypass peptide. In the graph, the mass to charge ratio (m/z) of each fragmented ion is on the x-axis and the percent of the base peak is shown on the y-axis. **(B)** RNA sequence of a portion of the Tm 9 sequence is shown as in **A**. The predicted amino acid of a region of the product of landing at GCA §45-47 is indicated above the RNA sequence. MS/MS analysis of the GCA §45-47 bypass peptide. In the graph, the mass to charge ratio (m/z) of each fragmented ion is on the x-axis and the percent of the base peak is shown on the y-axis. **(C)** RNA sequence of a portion of the Tm 9 sequence is shown as in **A**. The predicted amino acid of a region of the product of landing at GCA §48-50 is indicated above the RNA sequence. MS/MS analysis of the GCA §48-50 bypass peptide. In the graph, the mass to charge ratio (m/z) of each fragmented ion is on the x-axis and the percent of the base peak is shown on the y-axis. (The sequence of Tm9 and corresponding electrospray mass chromatogram is shown in Fig. 6.)
